# MRLocus: identifying causal genes mediating a trait through Bayesian estimation of allelic heterogeneity

**DOI:** 10.1101/2020.08.14.250720

**Authors:** Anqi Zhu, Nana Matoba, Emmaleigh Wilson, Amanda L. Tapia, Yun Li, Joseph G. Ibrahim, Jason L. Stein, Michael I. Love

## Abstract

Expression quantitative trait loci (eQTL) studies are used to understand the regulatory function of non-coding genome-wide association study (GWAS) risk loci, but colocalization alone does not demonstrate a causal relationship of gene expression affecting a trait. Evidence for mediation, that perturbation of gene expression in a given tissue or developmental context will induce a change in the downstream GWAS trait, can be provided by two-sample Mendelian Randomization (MR). Here, we introduce a new statistical method, MRLocus, for Bayesian estimation of the gene-to-trait effect from eQTL and GWAS summary data for loci displaying allelic heterogeneity, that is, containing multiple LD-independent eQTLs. MRLocus makes use of a colocalization step applied to each eQTL, followed by an MR analysis step across eQTLs. Additionally, our method involves estimation of allelic heterogeneity through a dispersion parameter, indicating variable mediation effects from each individual eQTL on the downstream trait. Our method is evaluated against state-of-the-art methods for estimation of the gene-to-trait mediation effect, using an existing simulation framework. In simulation, MRLocus often has the highest accuracy among competing methods, and in each case provides more accurate estimation of uncertainty as assessed through interval coverage. MRLocus is then applied to five causal candidate genes for mediation of particular GWAS traits, where gene-to-trait effects are concordant with those previously reported. We find that MRLocus’ estimation of the causal effect across eQTLs within a locus provides useful information for determining how perturbation of gene expression or individual regulatory elements will affect downstream traits. The MRLocus method is implemented as an R package available at https://mikelove.github.io/mrlocus.

## Introduction

Genome-wide association studies (GWAS) have identified many loci associated with complex traits and diseases. A major goal now is to understand the mechanism by which non-coding genetic variation influences trait levels through changes in gene expression. This involves identifying the causal variants at a locus, determining if the same variants are associated with both gene expression and trait, and disambiguating mediation from pleiotropy (Yao et al., 2020). Proposing mediating genes from existing expression quantitative trait loci (eQTL) and GWAS resources will lead to experiments that test whether modulating gene expression influences traits, and therefore inform further research and development of treatments.

Current efforts to identify the genes underlying GWAS risk often make use of either colocalization of GWAS signal with eQTLs, or expression imputation. In colocalization, statistical models are used to probabilistically assess if the same genetic variants within a locus are likely to be causally contributing to both eQTL and GWAS signals, taking into account the structure of linkage disequilibrium (LD) for a given population. A number of different methods have been proposed for colocalization, including QTLMatch (Plagnol et al., 2009), coloc (Giambartolomei et al., 2014; Wallace et al., 2012), eCAVIAR (Hormozdiari et al., 2016), enloc (Wen et al., 2017), RTC (Ongen et al., 2017), and Primo for multi-omics colocalization (Gleason et al., 2019). Expression imputation methods, as in transcriptome-wide association studies (TWAS), add additional information by including subthreshold signal for both GWAS and eQTL to identify which genes’ expression may have a non-zero local genetic correlation with a given GWAS trait (Gamazon et al., 2015; Gusev et al., 2016; Mancuso et al., 2019). Further refinements of TWAS statistical models have allowed for probabilistic fine-mapping within loci harboring multiple candidate genes by accounting for LD structure, as in FOCUS (Mancuso et al., 2019).

Though colocalization and expression imputation suggest genes involved in a trait, neither method is designed to disambiguate between pleiotropy and mediation. The latest generation of methods for determining those genes involved in mediating GWAS signal have combined the approaches of colocalization and expression imputation with statistical techniques from the field of Mendelian randomization (MR) (Davey Smith & Hemani, 2014; Smith & Ebrahim, 2003), as reviewed recently (Broekema et al., 2020). Intuitively, these methods work by determining if those genetic variants which influence gene expression also influence a downstream trait in proportionate degrees. Evidence for mediation is provided by randomized genetic variation used to perturb gene expression and observing the expression effects propagated to traits. With access to genotype, expression, and trait data, classical mediation techniques can be employed, as in the methods CIT (Millstein et al., 2009) and SMUT (Zhong et al., 2019), while MR-link (van der Graaf et al., 2019) makes use of individual-level data from an eQTL study and summary statistics from GWAS to perform MR analysis.

Other sets of methods testing gene-to-trait mediation require only summary statistics from eQTL and GWAS studies, as it may be prohibitive for a method to require access to the per-participant genotypes, expression values, and GWAS trait values (Pasaniuc & Price, 2017). Methods such as SMR (Summary data-based MR) consider genetic variants as potential instruments that may affect a downstream trait through the mediator of gene expression (Zhu et al., 2016). SMR uses the top cis-eQTL per gene for testing the gene-to-trait effect, and the authors proposed HEIDI (Heterogeneity In Dependent Instruments) to test for heterogeneity of the gene-to-trait effect across multiple variants in a region, with heterogeneity indicating that the gene may not be causally involved in the trait. CaMMEL performs causal mediation analysis across multiple loci, each harboring multiple candidate genes, and attempts to adjust for unmediated effects (Park et al., 2017). TWMR performs a multivariable MR analysis wherein the gene-to-trait effect of multiple genes is estimated simultaneously to reduce bias (Porcu et al., 2019). In contrast to SMR, TWMR includes multiple variants per gene for effect estimation. LDA-MR-Egger (Barfield et al., 2018) and PMR-Summary-Egger (Yuan et al., 2020) account for LD structure in estimating the gene-to-trait effect, and additionally allow for pleiotropy through the use of an intercept term. PTWAS (Zhang et al., 2019) adds to the analysis of gene-to-trait effects an upstream fine-mapping of cis-eQTL using DAP (Wen et al., 2016), and estimation of gene-to-trait effect heterogeneity in the case of multiple independent eQTL signal clusters, employing an I^2^ statistic (Higgins & Thompson, 2002). The I^2^ statistic ranges between 0 and 1 and represents, in the gene-to-trait meditation case, the percent of variance in estimated effect sizes across signal clusters that arises from true effect heterogeneity. MESC also attempts to determine whether gene expression mediates GWAS signals, and as with CaMMEL and TWMR, multiple genes are considered simultaneously (Yao et al., 2020). Finally, a new method MR-Robin performs robust MR with a focus on multiple tissue eQTL summary statistics, allowing for fewer and correlated candidate causal variants to achieve accurate gene-to-trait effect estimates (Gleason et al., 2020).

The ability to perform MR within a locus relies on having multiple independent “instruments”, SNPs which are found to be associated with the potential mediator, and which plausibly only affect the downstream trait through the mediator. Recently, it has been found that more than a third of genes have more than one independent cis-eQTL, with some genes having up to 13 independent cis-eQTL signals, detected by conditional analysis in peripheral blood (Jansen et al., 2017). A recent study integrating neonatal gene expression with GWAS of autoimmune and allergic disease performed MR analysis across 52 genes that had three or more cis-eQTLs (Huang et al., 2020). In addition, power to detect mediation at typical eQTL and GWAS sample sizes will depend on the percent of mediated heritability through a gene of interest. The MESC method estimated that the average percent of mediated heritability through the expression of *all* genes is around 11%, averaging over 42 GWAS diseases and traits, with the top mediated traits having around 30% mediated heritability through expression of all genes (Yao et al., 2020).

The existing methods for assessing whether expression of a particular gene in some context may mediate GWAS signal have primarily focused on their ability to perform genome-wide mediation scans across multiple tissues or cell types. This is a critical task in determining the genetic architecture and the most relevant molecular contexts for a trait (e.g. tissues, cell types, or developmental stage) which are often not known a priori. However, when considering functional follow-up experiments at an individual locus, investigators are interested in the degree to which modulation of gene expression results in trait differences. This can be quantified by the statistical uncertainty regarding a potential gene-to-trait effect, as well as the heterogeneity of gene-to-trait effects across independent signal clusters within a locus, as is the focus in PTWAS.

Here, we propose MRLocus, a Bayesian model for estimating the gene-to-trait effect from multiple independent signal clusters for one gene, as well as for estimating the heterogeneity of the effects across clusters. We have designed our method for prioritization of genes in functional experiments, where the genes under study have already been identified as candidates for mediation, having emerged from one of the global mediation scanning methods described above, or from colocalization or TWAS. MRLocus performs estimation of the gene-to-trait effect itself, as our focus is on experimental follow-up, as opposed to estimation of the percent of mediated heritability in a given population. In comparisons with other recently developed methods for identifying mediating genes from eQTL and GWAS summary data and LD matrices, TWMR and PTWAS, MRLocus was often more accurate in estimation of the gene-to-trait effect across simulated eQTL and GWAS experiments, and had higher and closer to nominal coverage of the true effect when considering its credible intervals. Using existing eQTL and GWAS data, we also demonstrate mediation is observed at previously reported and experimentally validated loci. The MRLocus method is implemented as an open source R package with full function documentation and a software vignette demonstrating its use, publicly available at https://mikelove.github.io/mrlocus.

## Methods

### MRLocus method

MRLocus consists of two steps, (1) colocalization and (2) MR slope fitting, each of which use Bayesian hierarchical models specified in the Stan probabilistic programming language, and with posterior inference using the Stan and RStan software packages (Carpenter et al., 2017). As in PTWAS (Zhang et al., 2019), MRLocus attempts to estimate the gene-to-trait effect or slope by identifying “LD-independent” signal clusters and then assesses the strength of evidence of mediation and the heterogeneity of the allelic effects. In referring to “LD-independent” clusters, we refer to non-overlapping sets of SNPs with low LD: MRLocus uses PLINK’s clumping algorithm (Purcell et al., 2007) to identify LD-independent sets of SNPs based on eQTL p-value (as discussed below), whereas PTWAS uses DAP (Wen et al., 2016) for identifying signal clusters. Alternatively, conditional analysis could be used as discussed in a recent coloc methods paper (Wallace, 2020). As with other gene mediation methods mentioned above, we focus here on common SNPs (using a minor allele frequency (MAF) filter on real and simulated data of 0.01). To the extent that a SNP or genetic variant gives rise to both the eQTL and GWAS signal and is in the set analyzed by MRLocus, then the model has a chance to find the “causal” SNP, although in general MRLocus may identify a SNP which is in high LD with the causal SNP.

In contrast to other methods for estimating gene-to-trait effects, MRLocus additionally performs a colocalization step prior to slope fitting, using eQTL and GWAS summary statistics (estimated coefficients and standard errors (SE)), based on LD matrices (either distinct matrices when eQTL/GWAS are performed in different populations, or a single shared matrix can be used when eQTL/GWAS are performed in the same population). The colocalization step attempts to identify a single candidate causal SNP per LD-independent signal cluster. Here “candidate causal SNP” refers to the hypothesis that the SNP gives rise to both the observed eQTL and GWAS signal, given the LD matrices. The colocalization step produces posterior estimates that assess the degree to which the summary statistics and LD matrices support the causal hypothesis per signal cluster (see Supplementary Methods for details on the statistical model). If the SNP is a strong candidate for causing both the eQTL and GWAS signal in a signal cluster, then the posterior estimates for the eQTL and GWAS effect sizes will be large (in absolute value) for the SNP, and near 0 for the other SNPs. If the SNP is a strong candidate for only the eQTL signal, but not for the GWAS signal, then the posterior estimate for the chosen SNP for the GWAS signal will be near 0. Finally, we note that prior to the colocalization step, MRLocus performs collapsing of highly correlated SNPs (threshold of 0.95 correlation), such that the final “per-SNP” results actually correspond to results for representatives from sets of highly correlated SNPs. MRLocus also performs allele flipping such that all SNPs are coded to be in positive LD correlation to the index SNP that has a positive estimated coefficient for eQTL (Supplementary Methods). This ensures that the statistical modeling and visualizations are always referring to the effect of an expression-increasing allele.

MRLocus’ colocalization step is motivated by the eCAVIAR model (Hormozdiari et al., 2016) as it formulates a generative model for the summary statistics based on true underlying signals, but is distinct from the eCAVIAR model in two respects. First, eCAVIAR models the *z*-scores from eQTL and GWAS, while MRLocus directly models the estimated coefficients, as our focus is on estimation of the gene-to-trait effect, which can be conceived as in other MR applications as a regression of coefficients from GWAS on eQTL. Second, eCAVIAR uses a multivariate normal distribution to model the vector of observed *z*-scores in a locus, while MRLocus uses a univariate distribution to model the estimated coefficients of the SNPs in each LD-independent signal cluster. The univariate distribution was chosen for its increased performance in accuracy and in efficiency in model fitting, as well as for flexibility in specification of prior distributions. In its implementation of colocalization, MRLocus uses a horseshoe prior (Carvalho et al., 2010) on the true coefficients for eQTL and GWAS signal per signal cluster, which helps to induce sparsity in the posterior estimates of the coefficients prior to mediation analysis (Supplementary Methods). The proposed use of the horseshoe prior during colocalization to identify putative causal SNPs from eQTL and GWAS coefficients within a signal cluster is distinct from other Bayesian MR methods’ use of the horseshoe prior on pleiotropic effects or on the mediation slope (Berzuini et al., 2020; Fazia et al., 2019; Uche-Ikonne et al., 2019).

MRLocus’ slope fitting step involves estimation of the gene-to-trait effect across signal clusters (Figure 1). For slope estimation, the best candidate SNP per LD-independent signal cluster is chosen, based on which SNP has the largest posterior mean of the eQTL effect size from the colocalization step. Again, a hierarchical model is used to perform inference on parameters of interest, in this case the slope (α) of true GWAS coefficients on true eQTL coefficients (Supplementary Methods). If the signal cluster does not provide evidence of colocalization, the estimate of the GWAS coefficient from the previous step will be near 0, and this will bring the estimated slope toward 0 as well. Quantile-based credible intervals on the slope coefficient provide information regarding the uncertainty of the gene mediating the trait measured in the GWAS. Finally, whereas PTWAS makes use of an I^2^ statistic (Higgins & Thompson, 2002) for quantifying the heterogeneity of the allelic effects at the locus, MRLocus estimates the dispersion (σ) of the different allelic effects around the predicted values given by the slope. Therefore, MRLocus may have high certainty on the slope (narrow credible interval for α not overlapping 0), while nevertheless estimating that the dispersion of allelic effects around the slope is large (σ). The tradeoff between uncertainty on the estimate of α and the dispersion σ of allelic effects around fitted line naturally depends upon the number of independent signal clusters at the locus. When a single signal cluster is provided by PLINK clumping, MRLocus uses a parametric bootstrap to estimate the gene-to-trait effect (Supplementary Methods).

**Figure 1:**
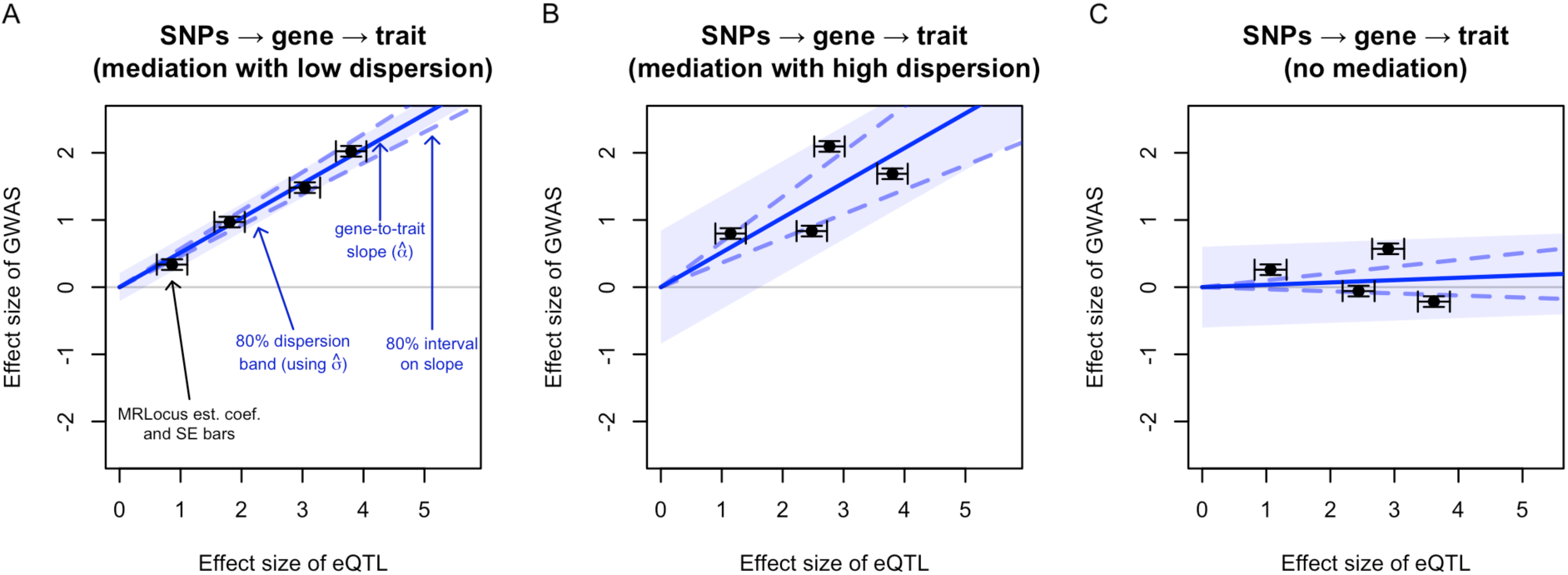
MRLocus estimates the gene-to-trait effect (solid blue line) as the slope from paired eQTL and GWAS effect sizes from independent signal clusters (black points with standard error bars), here on simulated coefficients. The dispersion of allelic effects around the main gene-to-trait effect (light blue band) is also estimated. An 80% credible interval on the slope is indicated with dashed blue lines sloping above and below the solid blue line. Panels represent loci demonstrating A) mediation with low dispersion, B) mediation with high dispersion, and C) colocalization of eQTL and GWAS signals but no evidence of mediation (slope credible interval overlaps 0). Investigators may wish to prioritize loci for experimental follow-up in which a typical “dosage” pattern is observed, such that alleles contributing small amounts to expression of a gene contribute small amounts to GWAS trait, and similarly for large effect alleles.

### Choice of methods for comparison

We chose to focus on TWMR (Porcu et al., 2019) and PTWAS (Zhang et al., 2019) in our comparisons, as these two methods had a focus on estimation of the gene-to-trait effect, were able to run on eQTL data for a single gene and a single tissue, and required only summary statistics and LD matrices. We additionally compared MRLocus to LDA-MR-Egger (Barfield et al., 2018) and PMR-Summary-Egger (Yuan et al., 2020) on the first simulation setting. Other methods that likewise determine if one or more genes may mediate traits include CaMMEL (Park et al., 2017), MESC (Yao et al., 2020), and MR-Robin (Gleason et al., 2020). We were not able to run CaMMEL using only LD matrices from the eQTL and/or GWAS cohort, as the fit.med.zqtl function takes genotype design matrices as input. We were not able to run MESC with less than 5 genes (our focus with MRLocus is on single gene mediating effect estimation). Finally, MR-Robin also provides robust estimates of gene-to-trait effects but with a focus on multiple-tissue eQTL summary statistics as input.

### Simulation

For simulation, we used the pre-existing TWAS simulation framework, twas_sim (mancusolab, n.d.), which simulates eQTL and GWAS datasets using real genotype data (1KG EUR Phase3) and outputs summary statistics. This software has a number of options for simulation parameters including *cis* heritability of gene expression (referred to here as “h^2^”), variance explained in the downstream trait by the gene’s expression level (referred to here as “var. exp.”), and the number of SNPs in a locus which are *cis* eQTL. A single gene’s expression was simulated per experiment, and both gene and trait were scaled to unit variance for slope estimation. Simulations of paired eQTL and GWAS datasets were performed where the gene was a partial mediator of the trait, as well as null simulations where the gene expression was unrelated to the downstream trait. TWMR, PTWAS, MRLocus, LDA-MR-Egger, and PMR-Summary-Egger were all run on the same summary data from eQTL and GWAS simulations. The Snakemake software (Koster & Rahmann, 2012) was used for automation of simulation scripts including specification of random seed for each of the 240 simulations, in order to assist with computational reproducibility of simulations. Simulation and analysis code is provided at https://github.com/mikelove/mrlocusPaper. The sample sizes for eQTL and GWAS were kept at their default values of N = 500 and N = 100,000, respectively. The percent of SNPs in a locus which are eQTLs was set to 1%. The gene h^2^ was varied from its default value (0.1) to a higher value (0.2) and a lower value (0.05), and the trait variance explained by gene expression was varied from its default value (0.01) to two lower values (0.005, 0.001, as well as to 0 indicating a null simulation where gene expression does not explain variation in a GWAS trait). 20 replications of each combination of 3 (for h^2^) × 4 (for variance explained) resulted in 240 simulations, 60 of which simulated no mediation of gene on trait (Supplementary Figure 1).

LD-based clumping implemented in PLINK (v1.90b) (Purcell et al., 2007) was performed on eQTL simulation un-adjusted p-values with the following settings: --clump-p1 0.001 --clump-p2 1 --clump-r2 0.2 --clump-kb 500. All PLINK clumps were provided to TWMR (commit 62994ec) (Porcu et al., 2019) and MRLocus (v0.0.14) for gene-to-trait effect estimation. PTWAS (v1.0) (Zhang et al., 2019) was provided with output from DAP (DAP-G, commit ac38301) (Wen et al., 2016) with settings: -d_n 500 -d_syy 500 (as twas_sim scales the GWAS trait to unit variance). LDA-MR-Egger (from <u>R script</u>) and PMR-Summary-Egger (v1.0) were supplied with the eQTL and GWAS summary statistics of the locus, without clumping. For all simulations, if there were no SNPs in the simulated locus with eQTL un-adjusted p-value < 0.001 then a new seed was drawn. For simulation comparisons, MR with inverse variance weighted (IVW) regression with fixed effects (Burgess et al., 2013) was computed using the true causal eQTLs (“causal”) or across all SNPs (“all”) using the mr_ivw_fe function in the TwoSampleMR R package (v0.5.5) (Hemani et al., 2018) with eQTL as the exposure study and GWAS as the outcome study. The “causal” IVW MR analysis served as an “oracle” estimator in the simulations, as it was provided with information not available in a typical analysis and not provided to other methods. The number of true eQTL SNPs, PLINK clumps, and DAP signal clusters are shown in Supplementary Figure 2. The distribution of the number of SNPs per clump before and after collapsing highly correlated SNPs (part of MRLocus pre-processing, described in Supplementary Methods) is shown in Supplementary Figure 3.

**Figure 2:**
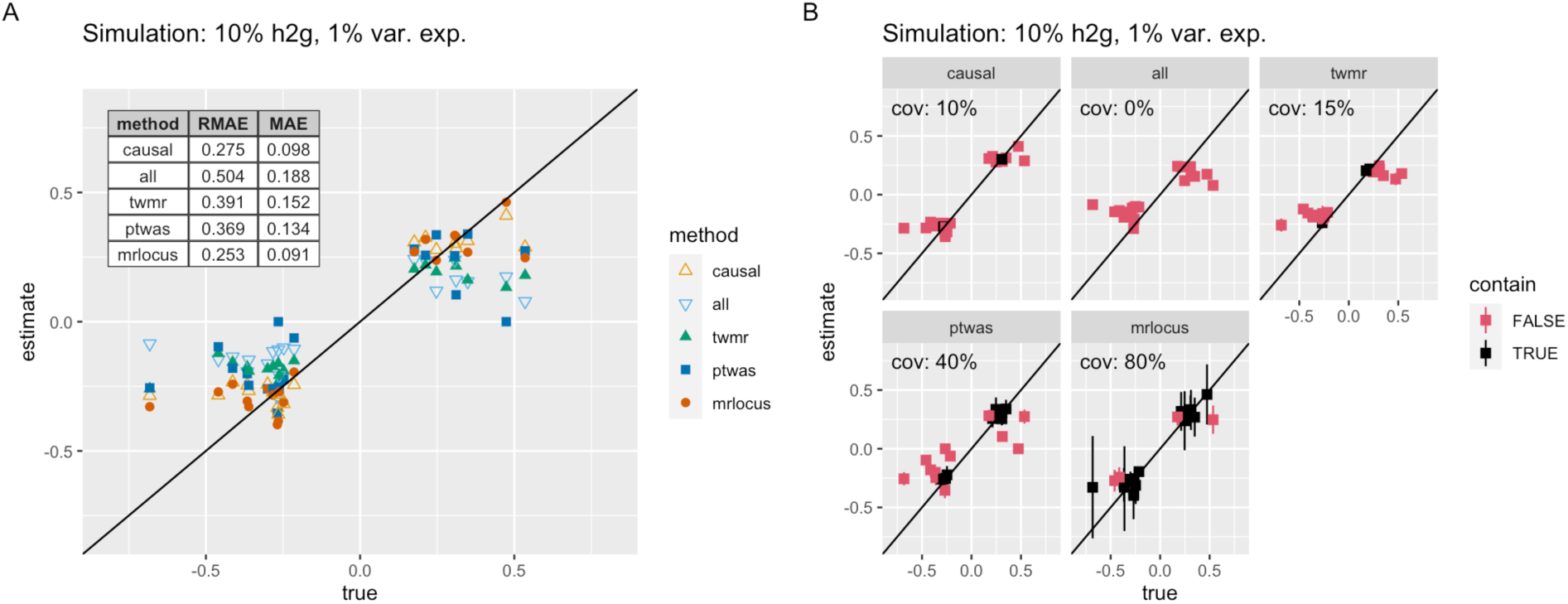
Performance of methods on 20 simulated eQTL and GWAS datasets. A) Estimates of each method over true (simulated) gene-to-trait values. The method denoted with “causal” indicates an inverse variance weighted slope estimation using the true causal SNPs but the estimated coefficients and SEs (an oracle estimate), and “all” indicates an inverse variance weighted slope estimation using all SNPs. B) Observed coverage (abbreviated cov. within each panel) of 80% confidence or credible intervals from each method. If the interval contains the true effect size, it is colored black, otherwise red.

**Figure 3:**
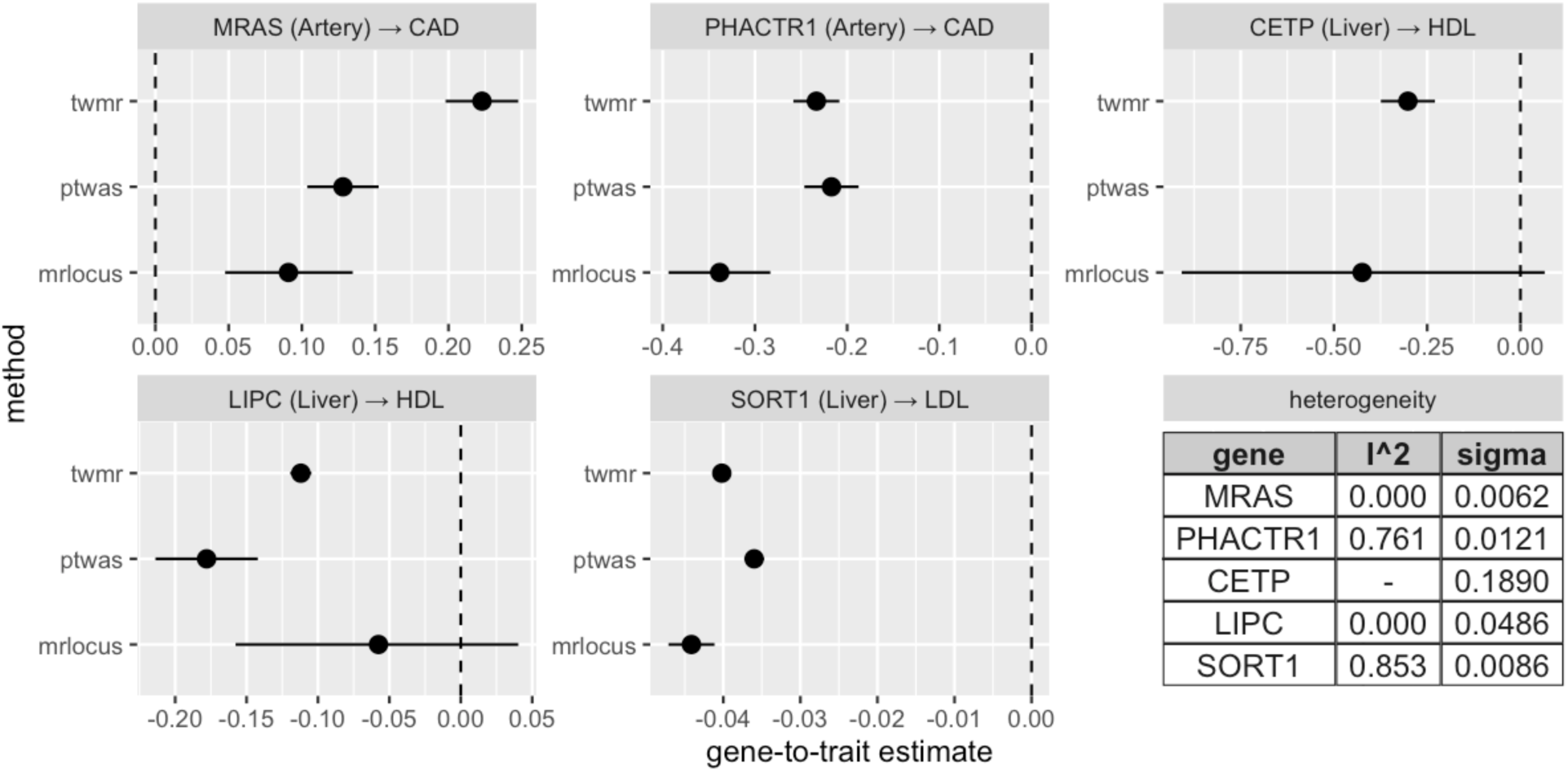
Estimated gene-to-trait effects for TWMR, PTWAS, and MRLocus on five eQTL and GWAS datasets for strong candidate genes for mediation of the GWAS trait. 80% confidence or credible intervals are shown (MRLocus provides quantile-based credible intervals). The artery eQTL were estimated from inverse normal transformed expression data from GTEx, while the liver meta-analysis eQTL were estimated from log2 expression data, thus the estimated slopes represent changes in SD of gene expression on log odds for CAD risk, and doubling of gene expression on lipid levels in population SD, respectively. The last panel provides two estimates of heterogeneity: PTWAS uses an I^2^ statistic to quantify heterogeneity across signal clusters (ranging from 0 to 1, for low to high heterogeneity), while MRLocus quantifies the dispersion with a parameter, s, on the scale of the estimated slope.

### Real data analysis

To evaluate the performance of methods on real data, we applied TWMR, PTWAS, and MRLocus to five eQTL-GWAS pairs chosen based on literature review of previous connections between gene expression and downstream phenotype: *MARS* (tibial artery) - coronary artery disease (CAD), *PHACTR1* (tibial artery) - CAD, *CETP* (liver) - high-density lipoproteins (HDL), *LIPC* (liver) - HDL, and *SORT1* (liver) - low-density lipoproteins (LDL). eQTL and GWAS summary data were obtained as summarized in Supplementary Tables 1 and 2 (SNP details and URLs). Briefly, eQTL data on tibial artery and liver was obtained from the Genotype-Tissue Expression (GTEx) project v8 (GTEx Consortium et al., 2017) and directly from the authors of a liver meta-analysis study (Strunz et al., 2018), respectively. GTEx v8 estimated eQTL coefficients were on the scale of unit variance expression values (following per-gene inverse normal transformation), while the liver meta-analysis effect sizes were on the scale of log2 normalized expression, according to references. GWAS summary statistics on CAD were obtained from CARDioGRAMplusC4D (Coronary ARtery DIsease Genome wide Replication and Meta-analysis (CARDIoGRAM) plus The Coronary Artery Disease (C4D) Genetics) consortium (Nikpay et al., 2015), where coefficients represent log odds ratios (OR), and the UK Biobank (https://www.ukbiobank.ac.uk) obtained from http://www.nealelab.is/uk-biobank/, where coefficients are estimated with respect to unit variance, continuous scale HDL or LDL.

Prior to defining independent signal clusters, we filtered out SNPs with MAF < 0.01 from GWAS data, and p-values were corrected with lambda GC (Bacanu et al., 2000). We used raw p-values for eQTL data and genomic control corrected p-values for GWAS data. As our task for downstream inference is estimation of the slope of GWAS coefficients over eQTL coefficients across signal clusters, it is not necessary that the eQTL signal clusters attain genome-wide significance before being provided to MRLocus.

### Defining independent signal clusters

We first identified LD-independent signal clusters for eQTL and GWAS datasets by LD-based clumping implemented in PLINK (1.90b) (Purcell et al., 2007) with the following settings: -- clump-p1 0.001 --clump-p2 1 --clump-kb 500 --r2 0.2 for eQTL, and --clump-p1 5e-8 --clump-p2 1 --clump-kb 500 --r2 0.2 for GWAS. LD (*r*^*2*^) was estimated in the European population from the 1000 Genome Project phase 3 (1KG EUR). Next, to define a candidate pair for colocalization, we calculated *r*^*2*^ between eQTL index SNPs and GWAS index SNPs for each cluster. The clusters are considered to be candidates for colocalization if *r*^*2*^ >= 0.4. The union of SNPs within paired clusters were provided to MRLocus.

We also kept the rest of the eQTL independent clusters, defined them as unpaired eQTL signals, and additionally provided these data to MRLocus. Because for those pairs, we do not have corresponding GWAS clusters, we directly obtained test statistics of matched SNPs from GWAS summary data. Unpaired eQTL signal clusters, which were not candidates for colocalization with GWAS, were included in all analyses, as these provide important evidence *against* the gene mediating the trait. However, as we are performing a unidirectional MR analysis of a candidate gene for mediation of downstream traits, unpaired GWAS signal clusters were not included.

### Generating MRLocus input files

For each independent pair, we generated MRLocus input files (effect size tables) with estimated coefficients, SE, and reference and effect alleles from both eQTL and GWAS. Pairwise LD (*r*) between SNPs metrics (LD metrics table) were generated by the plink --r function for all SNPs included in the corresponding effect size table using 1KG EUR. As LD matrix structure needs to be matched to the effect size table, the row of the effect size table was sorted by SNP position. We also included major/minor allele information from the reference panel (plink bim files) in the effect size table for proper allele flipping for statistical modeling and visualization (Supplementary Methods).

### Real data analysis with other methods

TWMR (Porcu et al., 2019) and PTWAS (Zhang et al., 2019) were used to compare estimates on real data loci. Both software packages require independent eQTL signal clusters and we used DAP (DAP-G, commit ac38301) (Wen et al., 2016) to estimate independent eQTL clusters for PTWAS (v1.0) as described in the original paper, while for TWMR (commit 62994ec) we ran PLINK clumping with the same parameters we used for MRLocus (v0.0.14) because we only have access to summary statistics (c.f. the original TWMR paper performed conditional analyses). DAP was applied to estimated coefficients and SE from eQTL summary statistics with default options except we set maximum models (-msize) to 20. PTWAS code was modified at <u>line 64</u> to allow for input of estimated coefficients and their SE, such that it provided slope estimates on the original scale of coefficients, not *z*-scores.

## Results

### Simulation

In order to evaluate the accuracy in estimating the mediated gene effect, we used a GWAS and eQTL simulation. The twas_sim simulation framework used for evaluating methods has default values for cis eQTL gene expression heritability of 10% (“h^2^”) and gene-mediated trait variance explained of 1% (“var. exp”). As the eQTL-based heritability could feasibly be higher or lower, we investigated values of 20% and 5% as well, which are within the range of detection for eQTL studies with hundreds of samples (Lloyd-Jones et al., 2017; Stranger et al., 2007). The default mediated trait variance explained value of 1% for a single gene is likely high, given that a recent publication using GTEx data has estimated the total gene-mediated heritability across all genes to be around 11% (±2%) averaging over 42 traits (Yao et al., 2020). We therefore considered even lower heritability of trait on gene expression of 0.5% and 0.1%, as well as a null simulation where gene expression did not mediate the downstream trait in any way (0%).

For the default values of 10% gene h^2^ and 1% trait variance explained, MRLocus had the highest accuracy in terms of relative mean absolute error (RMAE), dividing the error by the absolute value of the true slope, and in terms of mean absolute error (MAE) (Figure 2A). In addition, MRLocus was able to achieve nominal coverage for 80% credible intervals over the true values, while other methods had lower observed coverage, having too narrow confidence intervals (Figure 2B). We additionally tested two other methods, LDA-MR-Egger and PMR-Summary-Egger at the default twas_sim settings (Supplementary Figure 4). LDA-MR-Egger did not provide an estimate of SE for 6 of 20 simulations, while PMR did not provide an estimate of the effect for 14 of 20 simulations. These additional two methods had much higher error compared to TWMR, PTWAS, and MRLocus at the default twas_sim settings, and so we focused on the latter three methods for further evaluation. We note that in this first simulation, MRLocus obtained slightly lower RMAE of gene-to-trait slope compared to an oracle method that uses only the true causal SNPs (which are generally not known) and their estimated coefficients (“causal”) in IVW regression with fixed effects (Burgess et al., 2013). We believe the main reason for this improvement was MRLocus’ re-estimation of the coefficients in its colocalization procedure with shrinkage priors, resulting in coefficients that had lower variance potentially leading to lower error for the eQTL effect sizes (where N = 500).

**Figure 4:**
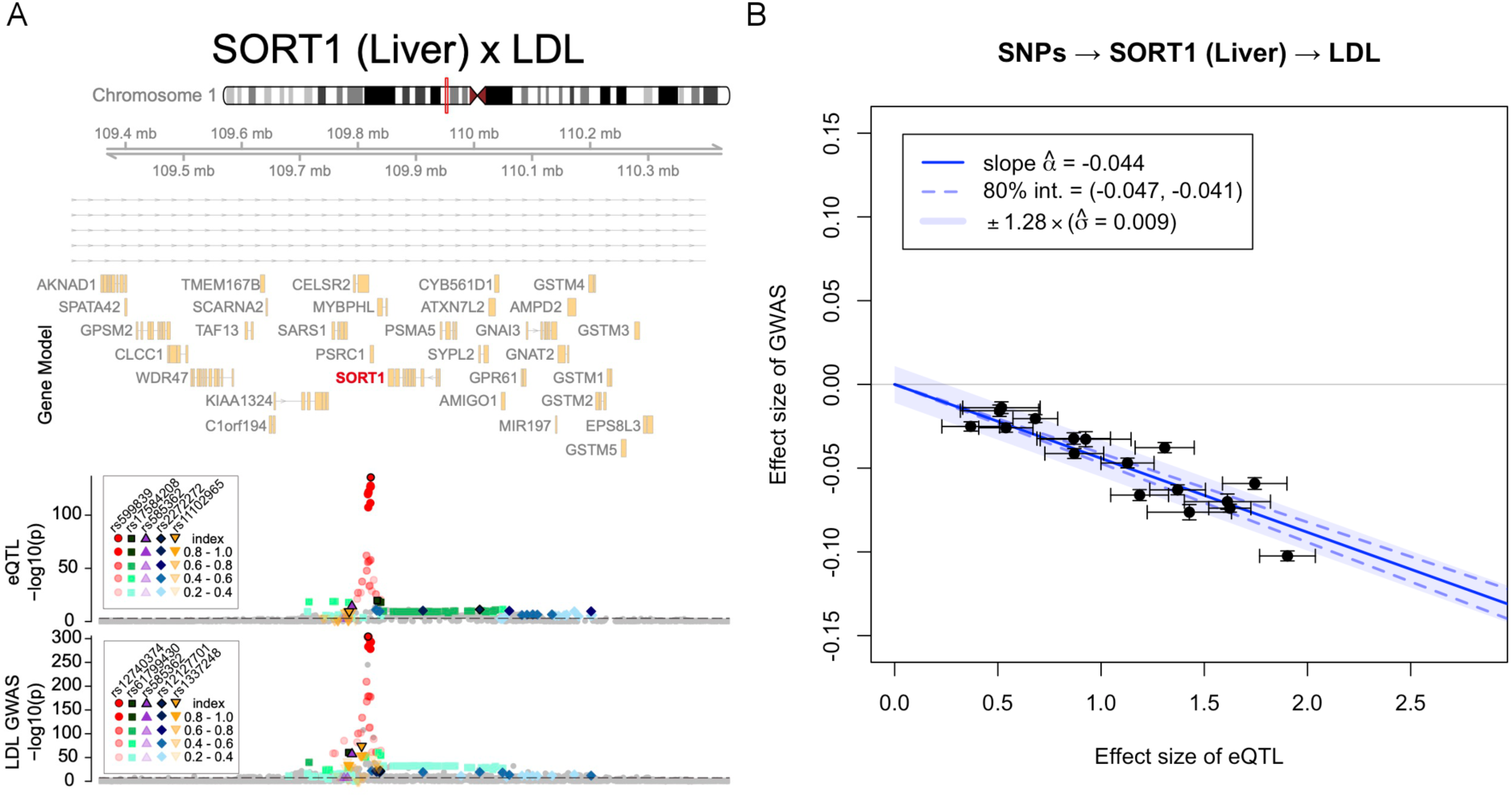
Colocalization and MRLocus estimation for *SORT1*. A) Colocalized signals in the *SORT1* region. From top panel to bottom, gene model (NCBI Refseq), eQTL for *SORT1* in liver (N = 588) (Strunz et al., 2018) and LDL association within UKBB (N = 343,621). LD was calculated to independent SNPs within 1KG EUR and colored accordingly. Symbols indicate independent co-localized (r^2^> 0.4) eQTL-GWAS pairs. Dashed line indicates a significance threshold at p = 0.001 or p = 5×10^−8^ for eQTL and GWAS respectively. B) MRLocus plot of the gene-to-trait effect for *SORT1* expression in liver on LDL levels. The signal clusters all provide consistent evidence for a gene-to-trait effect of −0.044, meaning that doubling of gene expression level in liver should reduce LDL by 4.4% of its population standard deviation. An 80% credible interval for the slope is indicated by dashed blue lines around the solid blue slope, while a range of heterogeneity of allelic effects is indicated by the light blue band.

For simulations using other parameter settings of h^2^ and gene-trait variance explained, the true causal SNPs with their estimated coefficients (“causal”) always had the lowest RMAE, as might be expected, but MRLocus had the second lowest RMAE in 4 of the remaining 8 non-null simulation settings with non-default values of h^2^ and variance explained (Supplementary Figure 5 – Supplementary Figure 12). For some simulations, PTWAS would report a division by a zero error had occurred, and in these cases, the estimate would not be included in PTWAS’ error or coverage calculations, and the estimate is plotted in figures at y=0. In the other 4 non-null settings, PTWAS had lower RMAE than MRLocus and higher than “causal”, although in 2 of these 4 settings PTWAS had division-by-zero errors for the majority of the replicate simulations, making comparisons between methods difficult. PTWAS and MRLocus always had lower RMAE than TWMR.

In terms of credible or confidence interval coverage, MRLocus always had better or equal coverage of the true values compared to all other methods within the non-null simulation settings, though it did not always reach the nominal level (Supplementary Figure 13 – Supplementary Figure 20). When gene h^2^ was 10% or 5% and trait variance explained was 1% or 0.5%, the nominal coverage (for 80% intervals) was achieved, with mean coverage of 79% over these 4 settings. When gene h^2^ was 20% (the highest setting) or when trait variance explained was 0.1% (the lowest setting), the coverage for MRLocus was lower than nominal, with mean coverage of 43%, though in all of these 5 settings MRLocus had better or equal interval coverage than the oracle method, TWMR, or PTWAS. The oracle method tended to have narrow confidence intervals, and MRLocus had higher coverage than the oracle method for all of the 9 non-null simulations. We believe the lower-than-nominal coverage seen here for the oracle method is likely from insufficient propagation of error during slope estimation. As the twas_sim framework does not include heterogeneity of effects from different signal clusters, an additional simulation was performed to assess MRLocus’ estimation of the dispersion of effects around the gene-to-trait fitted line (Supplementary Figure 21). Here, MRLocus was accurate both in estimation of the dispersion and quantification of uncertainty (attaining nominal credible interval coverage), with higher accuracy and smaller intervals as the number of LD-independent clusters increased, as expected.

In the 3 null simulation settings (gene h^2^ of 5%, 10%, 20% but no trait variance explained), MRLocus always had the highest coverage of the true slope value of 0, with average coverage of 95% (Supplementary Figure 22). Here, TWMR had average coverage of 65% and PTWAS had average coverage of 55%. The oracle method with the true eQTL SNPs had average coverage of 72%, and always lower than MRLocus. The oracle method again had narrow intervals as in the non-null simulations. As the oracle method uses GWAS SE for the true causal eSNPs for invariance variance weighting, we believe this was insufficient for propagation of error.

Across the 240 simulations, MRLocus was approximately 34 times slower than TWMR and 48 times slower than PTWAS, with a mean running time per locus of 121 seconds (compared to 3.5 seconds for TWMR and 2.5 seconds for PTWAS) (Supplementary Figure 23). This additional runtime is because MRLocus involves a colocalization step that is fit per signal cluster. Runtime for PLINK clumping and DAP were not included in the times presented above.

### Real data analysis

We compared TWMR, PTWAS, and MRLocus using eQTL and GWAS summary statistics for five gene-trait pairs in which there is strong evidence that the gene mediates the trait, and the direction of the effect has also been reported, such that we could compare our estimates against the literature. The paired index SNPs for the eQTL and GWAS datasets for these gene-trait pairs are provided in Supplementary Table 1, and LocusZoom-style plots (Hahne & Ivanek, 2016; Pruim et al., 2010) for the eQTL and GWAS tracks are provided in Supplementary Figure 24 – Supplementary Figure 27. Additionally, the LD patterns for paired PLINK clumps (eQTL and GWAS signals are candidates for colocalization) and unpaired PLINK clumps (eQTL only signals) are provided in Supplementary Figure 28 – Supplementary Figure 32.

On the five eQTL-GWAS dataset pairs, all three methods had consistent sign of the mediating effect (Figure 3), though MRLocus had larger credible intervals compared to confidence intervals from the other two methods, as was seen in the simulation datasets where MRLocus has improved coverage of the true effect size. In all cases, the sign of the mediating effect was in concordance with literature: higher *MRAS* (Artery) being hazardous for CAD (Alshahid et al., 2013; Song et al., 2019; Wu et al., 2015), higher *PHACTR1* (Artery) being protective for CAD (Chen et al., 2019; Codina-Fauteux et al., 2018), higher *CETP* (Liver) decreasing HDL levels (Tall, 2010), higher *LIPC* (Liver) decreasing HDL levels (Guerra et al., 1997), and higher *SORT1* (Liver) decreasing LDL levels (Musunuru et al., 2010; Strong et al., 2014; van der Graaf et al., 2019). For *CEPT* (Liver) paired with HDL, PTWAS encountered a division by zero error and did not produce an estimated effect. MRLocus had the strongest evidence for mediation with the *SORT1* (Liver) locus paired with LDL, with LocusZoom-style plot of the region in Figure 4A and MRLocus gene-to-trait estimate plot in Figure 4B. MRLocus plots for the remaining four gene-trait pairs are provided in Supplementary Figure 33.

On the 5 real data gene-trait pairs, MRLocus running with 2 cores had an average run time of 18.1 seconds per cluster, such that the average runtime of MRLocus per gene-trait pair was 3.1 minutes.

## Discussion

Here, we introduce MRLocus, a two-step Bayesian statistical procedure for estimation of gene-to-trait effects from eQTL and GWAS summary statistics. We find that MRLocus tends to have high accuracy in estimating the gene-to-trait effect across a variety of simulation settings, often higher than existing methods, and always had better credible interval coverage of true values, whether in mediating or null simulations. On real data analyses, MRLocus had consistent sign of estimates and comparable effect size compared to TWMR and PTWAS, but larger credible intervals compared to the other methods’ confidence intervals. While the effect sizes of alleles detected by GWAS on downstream traits examined here may be moderate, MRLocus’ estimation of the causal effect from perturbation of gene expression can be helpful in assessing the impact of therapeutic effects modulating expression on downstream traits (Visscher et al., 2017). For various systems, different downstream trait effect sizes qualify as practically meaningful increases or decreases, and MRLocus provides a framework for assessing what level of gene expression perturbation may be needed to obtain such changes in a downstream trait.

While the mediator evaluated by MRLocus in this work was gene expression effects via eQTL, the methods are generic, and protein abundance effects via pQTL could be used instead of eQTL. Two-sample MR linking pQTL and GWAS has already uncovered 30 metabolite features with evidence of causal effects on at least one disease (Qin et al., 2020), and a recent pQTL study of hepatic proteins reported a median of 4.5 local pQTL variants per protein (He et al., 2020), suggesting that there are loci with sufficient number of LD-independent clusters for MR analysis. Alternatively, pQTL could be used in place of the downstream GWAS trait in order to study mediation from gene expression to protein abundance (Buccitelli & Selbach, 2020), as previous work has found pQTL effect size to be positively correlated with eQTL effect size for variants ascertained through eQTL in human (Battle et al., 2015; Li et al., 2016), and that colocalized eQTL and pQTL signal leads to higher observed RNA-protein correlations in mice (Chick et al., 2016).

Given MRLocus’ improved performance with respect to interval coverage in the simulation, we feel that accurate estimation of uncertainty is an advantage to MRLocus, and the focus in developing a new method was on specificity for prioritization of gene targets for functional follow-up experiments. Additionally, the MRLocus model is extensible. The slope-fitting model could easily be generalized to use an alternative monotonic function, as long as there are sufficient LD-independent signal clusters to support fitting. On the other hand, MRLocus’ slower speed means it is likely not the best choice for a global scan of the transcriptome for mediating genes, while the other methods examined and discussed here have been successfully used to scan across all genes and across multiple tissues. Furthermore, MRLocus is designed for investigating loci with strong causal gene candidates, whereas other methods that estimate gene-to-trait effects for many genes in a locus simultaneously may have less biased estimation of the effect, when a strong gene candidate is not present. Future work on the MRLocus model may involve estimation of the mediating effect of candidate causal genes in the context of other relevant genes in a pathway and a polygenic background (Sinnott-Armstrong et al., 2020).

We note that our method focuses on common variation (MAF > 0.01), and that we collapse highly correlated SNPs to a single representative SNP during the pre-processing, such that we cannot determine if the final selected SNP is the true “causal” SNP. Future development of MRLocus could involve upstream use of methods defining credible sets (Hormozdiari et al., 2014; Hutchinson et al., 2020; Kichaev et al., 2014; Servin & Stephens, 2005; Wang et al., 2020; Wellcome Trust Case Control Consortium et al., 2012) or modeling based on a posterior inclusion probability as in LLARRMA or DAP (Valdar et al., 2012; Wen et al., 2016). Therefore, the current implementation of MRLocus can perform fine-mapping to the level of a highly correlated set of SNPs, which may be sufficient for identifying one or more regulatory elements (RE) to prioritize for functional follow-up experiments. The current implementation of MRLocus assumes that the mediation slope passes through the origin, and therefore that eQTL signal clusters do not affect the downstream trait through genes other than the eGene. Further iterations of MRLocus could relax this assumption through the addition of an intercept term accounting for invalid instruments as in MR-Egger (Bowden et al., 2015; Burgess et al., 2017). Finally, complementary information linking RE to genes, e.g. as measured by Hi-C, was not examined here, but have been proven successful elsewhere (HUGIN (Martin et al., 2017), H-MAGMA (Sey et al., 2020)), and we envision that prioritization of signal clusters in MRLocus that are supported by Hi-C would increase its power to detect causal genes.

As part of the MR analysis, MRLocus provides an estimate of the dispersion of effects around the estimated slope from LD-independent signal clusters, analogous to PTWAS’ use of the I^2^ statistic for effect size heterogeneity. The combined information from PTWAS and MRLocus regarding both uncertainty in estimation of the gene-to-trait slope, and estimated dispersion or heterogeneity of effects is critical when modeling context-specific (e.g. relevant tissue, cell type, or developmental stage) gene expression as a mediator for downstream traits. Different combinations of eQTL and GWAS SE (primarily influenced by sample size), extent of heterogeneity of effects, and the number of LD-independent signal clusters within a locus all may give rise to the same gene-to-trait effect and SE, but disentangling these sources of variance is important for experimental planning. For example, consider experimental follow-up for endophenotype downstream traits that could be feasibly measured *in vitro*. An investigator could choose between modulating gene expression directly or modulating the activity of an RE harboring candidate causal SNPs. A nonzero gene-to-trait effect with narrow credible interval estimated by MRLocus (as in Figure 1A and Figure 4B) would suggest that modulating the gene should affect the downstream trait, and the predicted effect could be assessed experimentally. However, high dispersion around the gene-to-trait slope (as in Figure 1B) suggests that perturbation of an RE implicated by candidate causal SNPs may induce an effect on the trait that is far from the effect size indicated by the fitted line. MRLocus provides a band around the predominant gene-to-trait slope, such that functional experiments per RE can therefore be prioritized. In all, here we demonstrate the MRLocus method and software utilizing summary statistics from eQTL and GWAS to identify genes mediating traits that can be prioritized for experimental follow-up.

## Acknowledgments

The authors would like to acknowledge the following individuals for helpful comments and suggestions on the work: Karen Mohlke, Laura Raffield, Robert Gentleman, and Steven Munger. The authors would like to thank Nicholas Mancuso for assistance in maintaining the ‘twas_sim’ simulation framework. The authors would like to thank Tobias Strunz and Bernhard Weber for their help in providing summary statistics for the three genes from their liver meta-analysis eQTL study.

The Genotype-Tissue Expression (GTEx) Project was supported by the <u>Common Fund</u> of the Office of the Director of the National Institutes of Health, and by NCI, NHGRI, NHLBI, NIDA, NIMH, and NINDS.

## References

Alshahid, M., Wakil, S. M., Al-Najai, M., Muiya, N. P., Elhawari, S., Gueco, D., Andres, E., Hagos, S., Mazhar, N., Meyer, B. F., & Dzimiri, N. (2013). New susceptibility locus for obesity and dyslipidaemia on chromosome 3q22.3. Human Genomics, 7, 15.

Bacanu, S. A., Devlin, B., & Roeder, K. (2000). The power of genomic control. American Journal of Human Genetics, 66(6), 1933–1944.

Barfield, R., Feng, H., Gusev, A., Wu, L., Zheng, W., Pasaniuc, B., & Kraft, P. (2018). Transcriptome-wide association studies accounting for colocalization using Egger regression. Genetic Epidemiology, 42(5), 418–433.

Battle, A., Khan, Z., Wang, S. H., Mitrano, A., Ford, M. J., Pritchard, J. K., & Gilad, Y. (2015). Genomic variation. Impact of regulatory variation from RNA to protein. Science, 347(6222), 664–667.

Berzuini, C., Guo, H., Burgess, S., & Bernardinelli, L. (2020). A Bayesian approach to Mendelian randomization with multiple pleiotropic variants. Biostatistics, 21(1), 86–101.

Bowden, J., Davey Smith, G., & Burgess, S. (2015). Mendelian randomization with invalid instruments: effect estimation and bias detection through Egger regression. International Journal of Epidemiology, 44(2), 512–525.

Broekema, R. V., Bakker, O. B., & Jonkers, I. H. (2020). A practical view of fine-mapping and gene prioritization in the post-genome-wide association era. Open Biology, 10(1), 190221.

Buccitelli, C., & Selbach, M. (2020). mRNAs, proteins and the emerging principles of gene expression control. Nature Reviews. Genetics. https://doi.org/10.1038/s41576-020-0258-4

Burgess, S., Bowden, J., Fall, T., Ingelsson, E., & Thompson, S. G. (2017). Sensitivity Analyses for Robust Causal Inference from Mendelian Randomization Analyses with Multiple Genetic Variants. Epidemiology, 28(1), 30–42.

Burgess, S., Butterworth, A., & Thompson, S. G. (2013). Mendelian randomization analysis with multiple genetic variants using summarized data. Genetic Epidemiology, 37(7), 658–665.

Carpenter, B., Gelman, A., Hoffman, M. D., Lee, D., Goodrich, B., Betancourt, M., Brubaker, M., Guo, J., Li, P., & Riddell, A. (2017). Stan : A Probabilistic Programming Language. Journal of Statistical Software, 76(1). https://doi.org/10.18637/jss.v076.i01

Carvalho, C. M., Polson, N. G., & Scott, J. G. (2010). The horseshoe estimator for sparse signals. In Biometrika (vol. 97, Issue 2, pp. 465–480). https://doi.org/10.1093/biomet/asq017

Chen, L., Qian, H., Luo, Z., Li, D., Xu, H., Chen, J., He, P., Zhou, X., Zhang, T., Chen, J., & Min, I. (2019). PHACTR1 gene polymorphism with the risk of coronary artery disease in Chinese Han population. Postgraduate Medical Journal, 95(1120), 67–71.

Chick, J. M., Munger, S. C., Simecek, P., Huttlin, E. L., Choi, K., Gatti, D. M., Raghupathy, N., Svenson, K. L., Churchill, G. A., & Gygi, S. P. (2016). Defining the consequences of genetic variation on a proteome-wide scale. Nature, 534(7608), 500–505.

Codina-Fauteux, V.-A., Beaudoin, M., Lalonde, S., Lo, K. S., & Lettre, G. (2018). PHACTR1 splicing isoforms and eQTLs in atherosclerosis-relevant human cells. BMC Medical Genetics, 19(1), 97.

Davey Smith, G., & Hemani, G. (2014). Mendelian randomization: genetic anchors for causal inference in epidemiological studies. Human Molecular Genetics, 23(R1), R89–R98.

Fazia, T., Egidi, L., Ayoglu, B., Beecham, A., Bitti, P. P., Ticca, A., McCauley, J. L., Nilsson, P., Berzuini, C., & Bernardinelli, L. (2019). Bayesian Mendelian Randomization identifies disease causing proteins via pedigree data, partially observed exposures and correlated instruments. http://arxiv.org/abs/1903.00682

Gamazon, E. R., Wheeler, H. E., Shah, K. P., Mozaffari, S. V., Aquino-Michaels, K., Carroll, R. J., Eyler, A. E., Denny, J. C., GTEx Consortium, Nicolae, D. L., Cox, N. J., & Im, H. K. (2015). A gene-based association method for mapping traits using reference transcriptome data. Nature Genetics, 47(9), 1091–1098.

Giambartolomei, C., Vukcevic, D., Schadt, E. E., Franke, L., Hingorani, A. D., Wallace, C., & Plagnol, V. (2014). Bayesian test for colocalisation between pairs of genetic association studies using summary statistics. PLoS Genetics, 10(5), e1004383.

Gleason, K. J., Yang, F., & Chen, L. S. (2020). A robust two-sample Mendelian Randomization method integrating GWAS with multi-tissue eQTL summary statistics. In Genetics (No. biorxiv;2020.06.04.135541v1; p. e1007889). bioRxiv.

Gleason, K. J., Yang, F., Pierce, B. L., He, X., & Chen, L. S. (2019). Primo: integration of multiple GWAS and omics QTL summary statistics for elucidation of molecular mechanisms of trait-associated SNPs and detection of pleiotropy in complex traits. In Genomics (No. biorxiv;579581v3; p. 1825). bioRxiv.

GTEx Consortium, Laboratory, Data Analysis &Coordinating Center (LDACC)—Analysis Working Group, Statistical Methods groups—Analysis Working Group, Enhancing GTEx (eGTEx) groups, NIH Common Fund, NIH/NCI, NIH/NHGRI, NIH/NIMH, NIH/NIDA, Biospecimen Collection Source Site—NDRI, Biospecimen Collection Source Site—RPCI, Biospecimen Core Resource—VARI, Brain Bank Repository—University of Miami Brain Endowment Bank, Leidos Biomedical—Project Management, ELSI Study, Genome Browser Data Integration &Visualization—EBI, Genome Browser Data Integration & Visualization—UCSC Genomics Institute, University of California Santa Cruz, Lead analysts:, Laboratory, Data Analysis &Coordinating Center (LDACC):, … Montgomery. (2017). Genetic effects on gene expression across human tissues. Nature, 550(7675), 204–213.

Guerra, R., Wang, J., Grundy, S. M., & Cohen, J. C. (1997). A hepatic lipase (LIPC) allele associated with high plasma concentrations of high density lipoprotein cholesterol. Proceedings of the National Academy of Sciences of the United States of America, 94(9), 4532–4537.

Gusev, A., Ko, A., Shi, H., Bhatia, G., Chung, W., Penninx, B. W. J. H., Jansen, R., de Geus, E. J. C., Boomsma, D. I., Wright, F. A., Sullivan, P. F., Nikkola, E., Alvarez, M., Civelek, M., Lusis, A. J., Lehtimäki, T., Raitoharju, E., Kähönen, M., Seppälä, I., … Pasaniuc, B. (2016). Integrative approaches for large-scale transcriptome-wide association studies. Nature Genetics, 48(3), 245–252.

Hahne, F., & Ivanek, R. (2016). Visualizing Genomic Data Using Gviz and Bioconductor. In E. Mathé & S. Davis (Eds.), Statistical Genomics (vol. 1418, pp. 335–351). Springer New York.

He, B., Shi, J., Wang, X., Jiang, H., & Zhu, H.-J. (2020). Genome-wide pQTL analysis of protein expression regulatory networks in the human liver. BMC Biology, 18(1), 97.

Hemani, G., Zheng, J., Elsworth, B., Wade, K. H., Haberland, V., Baird, D., Laurin, C., Burgess, S., Bowden, J., Langdon, R., Tan, V. Y., Yarmolinsky, J., Shihab, H. A., Timpson, N. J., Evans, D. M., Relton, C., Martin, R. M., Davey Smith, G., Gaunt, T. R., & Haycock, P. C. (2018). The MR-Base platform supports systematic causal inference across the human phenome. eLife, 7. https://doi.org/10.7554/eLife.34408

Higgins, J. P. T., & Thompson, S. G. (2002). Quantifying heterogeneity in a meta-analysis. In Statistics in Medicine (vol. 21, Issue 11, pp. 1539–1558). https://doi.org/10.1002/sim.1186

Hormozdiari, F., Kostem, E., Kang, E. Y., Pasaniuc, B., & Eskin, E. (2014). Identifying causal variants at loci with multiple signals of association. Genetics, 198(2), 497–508.

Hormozdiari, F., van de Bunt, M., Segrè, A. V., Li, X., Joo, J. W. J., Bilow, M., Sul, J. H., Sankararaman, S., Pasaniuc, B., & Eskin, E. (2016). Colocalization of GWAS and eQTL Signals Detects Target Genes. American Journal of Human Genetics, 99(6), 1245–1260.

Huang, Q. Q., Tang, H. H. F., Teo, S. M., Mok, D., Ritchie, S. C., Nath, A. P., Brozynska, M., Salim, A., Bakshi, A., Holt, B. J., Khor, C. C., Sly, P. D., Holt, P. G., Holt, K. E., & Inouye, M. (2020). Neonatal genetics of gene expression reveal potential origins of autoimmune and allergic disease risk. Nature Communications, 11(1), 3761.

Hutchinson, A., Watson, H., & Wallace, C. (2020). Improving the coverage of credible sets in Bayesian genetic fine-mapping. PLoS Computational Biology, 16(4), e1007829.

Jansen, R., Hottenga, J.-J., Nivard, M. G., Abdellaoui, A., Laport, B., de Geus, E. J., Wright, F. A., Penninx, B. W. J. H., & Boomsma, D. I. (2017). Conditional eQTL analysis reveals allelic heterogeneity of gene expression. Human Molecular Genetics, 26(8), 1444–1451.

Kichaev, G., Yang, W.-Y., Lindstrom, S., Hormozdiari, F., Eskin, E., Price, A. L., Kraft, P., & Pasaniuc, B. (2014). Integrating functional data to prioritize causal variants in statistical fine-mapping studies. PLoS Genetics, 10(10), e1004722.

Koster, J., & Rahmann, S. (2012). Snakemake--a scalable bioinformatics workflow engine. In Bioinformatics (vol. 28, Issue 19, pp. 2520–2522). https://doi.org/10.1093/bioinformatics/bts480

Li, Y. I., van de Geijn, B., Raj, A., Knowles, D. A., Petti, A. A., Golan, D., Gilad, Y., & Pritchard, J. K. (2016). RNA splicing is a primary link between genetic variation and disease. Science, 352(6285), 600–604.

Lloyd-Jones, L. R., Holloway, A., McRae, A., Yang, J., Small, K., Zhao, J., Zeng, B., Bakshi, A., Metspalu, A., Dermitzakis, M., Gibson, G., Spector, T., Montgomery, G., Esko, T., Visscher, P. M., & Powell, J. E. (2017). The Genetic Architecture of Gene Expression in Peripheral Blood. American Journal of Human Genetics, 100(2), 228–237.

mancusolab. (n.d.). mancusolab/twas_sim. Retrieved August 11, 2020, from https://github.com/mancusolab/twas_sim

Mancuso, N., Freund, M. K., Johnson, R., Shi, H., Kichaev, G., Gusev, A., & Pasaniuc, B. (2019). Probabilistic fine-mapping of transcriptome-wide association studies. Nature Genetics, 51(4), 675–682.

Martin, J. S., Xu, Z., Reiner, A. P., Mohlke, K. L., Sullivan, P., Ren, B., Hu, M., & Li, Y. (2017). HUGIn: Hi-C Unifying Genomic Interrogator. Bioinformatics, 33(23), 3793–3795.

Millstein, J., Zhang, B., Zhu, J., & Schadt, E. E. (2009). Disentangling molecular relationships with a causal inference test. BMC Genetics, 10, 23.

Musunuru, K., Strong, A., Frank-Kamenetsky, M., Lee, N. E., Ahfeldt, T., Sachs, K. V., Li, X., Li, H., Kuperwasser, N., Ruda, V. M., Pirruccello, J. P., Muchmore, B., Prokunina-Olsson, L., Hall, J. L., Schadt, E. E., Morales, C. R., Lund-Katz, S., Phillips, M. C., Wong, J., … Rader, D. J. (2010). From noncoding variant to phenotype via SORT1 at the 1p13 cholesterol locus. Nature, 466(7307), 714–719.

Nikpay, M., Goel, A., Won, H.-H., Hall, L. M., Willenborg, C., Kanoni, S., Saleheen, D., Kyriakou, T., Nelson, C. P., Hopewell, J. C., & Others. (2015). A comprehensive 1000 Genomes--based genome-wide association meta-analysis of coronary artery disease. Nature Genetics, 47(10), 1121.

Ongen, H., Brown, A. A., Delaneau, O., Panousis, N. I., Nica, A. C., & Dermitzakis, E. T. (2017). Estimating the causal tissues for complex traits and diseases. Nature Genetics, 49(12), 1676–1683.

Park, Y., Sarkar, A. K., He, L., Davila-Velderrain, J., De Jager, P. L., & Kellis, M. (2017). A Bayesian approach to mediation analysis predicts 206 causal target genes in Alzheimer’s disease. In Genetics (No. biorxiv;219428v3; p. 353). bioRxiv.

Pasaniuc, B., & Price, A. L. (2017). Dissecting the genetics of complex traits using summary association statistics. Nature Reviews. Genetics, 18(2), 117–127.

Plagnol, V., Smyth, D. J., Todd, J. A., & Clayton, D. G. (2009). Statistical independence of the colocalized association signals for type 1 diabetes and RPS26 gene expression on chromosome 12q13. Biostatistics, 10(2), 327–334.

Porcu, E., Rüeger, S., Lepik, K., eQTLGen Consortium, BIOS Consortium, Santoni, F. A., Reymond, A., & Kutalik, Z. (2019). Mendelian randomization integrating GWAS and eQTL data reveals genetic determinants of complex and clinical traits. Nature Communications, 10(1), 3300.

Pruim, R. J., Welch, R. P., Sanna, S., Teslovich, T. M., Chines, P. S., Gliedt, T. P., Boehnke, M., Abecasis, G. R., & Willer, C. J. (2010). LocusZoom: regional visualization of genome-wide association scan results. Bioinformatics, 26(18), 2336–2337.

Purcell, S., Neale, B., Todd-Brown, K., Thomas, L., Ferreira, M. A. R., Bender, D., Maller, J., Sklar, P., de Bakker, P. I. W., Daly, M. J., & Sham, P. C. (2007). PLINK: a tool set for whole-genome association and population-based linkage analyses. American Journal of Human Genetics, 81(3), 559–575.

Qin, Y., Meric, G., Long, T., Watrous, J., Burgess, S., Havulinna, A., Ritchie, S. C., Brozynska, M., Jousilahti, P., Perola, M., Lahti, L., Niiranen, T., Cheng, S., Salomaa, V., Jain, M., & Inouye, M. (2020). Genome-wide association and Mendelian randomization analysis prioritizes bioactive metabolites with putative causal effects on common diseases. In Genetic and Genomic Medicine (No. medrxiv;2020.08.01.20166413v1). medRxiv.

Servin, B., & Stephens, M. (2005). Imputation-based analysis of association studies: candidate regions and quantitative traits. In PLoS Genetics: Vol. preprint (Issue 2007, p. e114). https://doi.org/10.1371/journal.pgen.0030114.eor

Sey, N. Y. A., Hu, B., Mah, W., Fauni, H., McAfee, J. C., Rajarajan, P., Brennand, K. J., Akbarian, S., & Won, H. (2020). A computational tool (H-MAGMA) for improved prediction of brain-disorder risk genes by incorporating brain chromatin interaction profiles. Nature Neuroscience, 23(4), 583–593.

Sinnott-Armstrong, N., Naqvi, S., Rivas, M., & Pritchard, J. K. (2020). GWAS of three molecular traits highlights core genes and pathways alongside a highly polygenic background. In Genetics (No. biorxiv;2020.04.20.051631v1; p. 352). bioRxiv.

Smith, G. D., & Ebrahim, S. (2003). “Mendelian randomization”: can genetic epidemiology contribute to understanding environmental determinants of disease? International Journal of Epidemiology, 32(1), 1–22.

Song, Y., Ma, R., & Zhang, H. (2019). The influence of MRAS gene variants on ischemic stroke and serum lipid levels in Chinese Han population. Medicine, 98(48), e18065.

Stranger, B. E., Nica, A. C., Forrest, M. S., Dimas, A., Bird, C. P., Beazley, C., Ingle, C. E., Dunning, M., Flicek, P., Koller, D., Montgomery, S., Tavaré, S., Deloukas, P., & Dermitzakis, E. T. (2007). Population genomics of human gene expression. Nature Genetics, 39(10), 1217–1224.

Strong, A., Patel, K., & Rader, D. J. (2014). Sortilin and lipoprotein metabolism: making sense out of complexity. Current Opinion in Lipidology, 25(5), 350–357.

Strunz, T., Grassmann, F., Gayán, J., Nahkuri, S., Souza-Costa, D., Maugeais, C., Fauser, S., Nogoceke, E., & Weber, B. H. F. (2018). A mega-analysis of expression quantitative trait loci (eQTL) provides insight into the regulatory architecture of gene expression variation in liver. Scientific Reports, 8(1), 5865.

Tall, A. R. (2010). Functions of cholesterol ester transfer protein and relationship to coronary artery disease risk. Journal of Clinical Lipidology, 4(5), 389–393.

Uche-Ikonne, O. O., Dondelinger, F., & Palmer, T. (2019). Bayesian estimation of IVW and MR-Egger models for two-sample Mendelian randomization studies. In Epidemiology (No. medrxiv;19005868v1). medRxiv.

Valdar, W., Sabourin, J., Nobel, A., & Holmes, C. C. (2012). Reprioritizing genetic associations in hit regions using LASSO-based resample model averaging. Genetic Epidemiology, 36(5), 451–462.

van der Graaf, A., Claringbould, A., Rimbert, A., BIOS consortium, Westra, H.-J., Li, Y., Wijmenga, C., & Sanna, S. (2019). A novel Mendelian randomization method identifies causal relationships between gene expression and low-density lipoprotein cholesterol levels. In Genetics (No. biorxiv;671537v1; p. 303). bioRxiv.

Visscher, P. M., Wray, N. R., Zhang, Q., Sklar, P., McCarthy, M. I., Brown, M. A., & Yang, J. (2017). 10 Years of GWAS Discovery: Biology, Function, and Translation. American Journal of Human Genetics, 101(1), 5–22.

Wallace, C. (2020). Eliciting priors and relaxing the single causal variant assumption in colocalisation analyses. PLoS Genetics, 16(4), e1008720.

Wallace, C., Rotival, M., Cooper, J. D., Rice, C. M., Yang, J. H. M., McNeill, M., Smyth, D. J., Niblett, D., Cambien, F., Cardiogenics Consortium, Tiret, L., Todd, J. A., Clayton, D. G., & Blankenberg, S. (2012). Statistical colocalization of monocyte gene expression and genetic risk variants for type 1 diabetes. Human Molecular Genetics, 21(12), 2815–2824.

Wang, G., Sarkar, A., Carbonetto, P., & Stephens, M. (2020). A simple new approach to variable selection in regression, with application to genetic fine mapping. Journal of the Royal Statistical Society. Series B, Statistical Methodology, 25, 1.

Wellcome Trust Case Control Consortium, Maller, J. B., McVean, G., Byrnes, J., Vukcevic, D., Palin, K., Su, Z., Howson, J. M. M., Auton, A., Myers, S., Morris, A., Pirinen, M., Brown, M. A., Burton, P. R., Caulfield, M. J., Compston, A., Farrall, M., Hall, A. S., Hattersley, A. T., … Donnelly, P. (2012). Bayesian refinement of association signals for 14 loci in 3 common diseases. Nature Genetics, 44(12), 1294–1301.

Wen, X., Lee, Y., Luca, F., & Pique-Regi, R. (2016). Efficient Integrative Multi-SNP Association Analysis via Deterministic Approximation of Posteriors. American Journal of Human Genetics, 98(6), 1114–1129.

Wen, X., Pique-Regi, R., & Luca, F. (2017). Integrating molecular QTL data into genome-wide genetic association analysis: Probabilistic assessment of enrichment and colocalization. PLoS Genetics, 13(3), e1006646.

Wu, J., Yin, R.-X., Guo, T., Lin, Q.-Z., Shi, G.-Y., Sun, J.-Q., Shen, S.-W., Wang, Y.-M., Li, H., & Wu, J.-Z. (2015). Association between the MARS rs6782181 polymorphism and serum lipid levels. International Journal of Clinical and Experimental Pathology, 8(2), 1855–1866.

Yao, D. W., O’Connor, L. J., Price, A. L., & Gusev, A. (2020). Quantifying genetic effects on disease mediated by assayed gene expression levels. Nature Genetics, 52(6), 626–633.

Yuan, Z., Zhu, H., Zeng, P., Yang, S., Sun, S., Yang, C., Liu, J., & Zhou, X. (2020). Testing and controlling for horizontal pleiotropy with probabilistic Mendelian randomization in transcriptome-wide association studies. Nature Communications, 11(1), 3861.

Zhang, Y., Quick, C., Yu, K., Barbeira, A., The GTEx Consortium, Luca, F., Pique-Regi, R., Im, H. K., & Wen, X. (2019). Investigating tissue-relevant causal molecular mechanisms of complex traits using probabilistic TWAS analysis. In bioRxiv (p. 808295). https://doi.org/10.1101/808295

Zhong, W., Spracklen, C. N., Mohlke, K. L., Zheng, X., Fine, J., & Li, Y. (2019). Multi-SNP mediation intersection-union test. Bioinformatics, 35(22), 4724–4729.

Zhu, Z., Zhang, F., Hu, H., Bakshi, A., Robinson, M. R., Powell, J. E., Montgomery, G. W., Goddard, M. E., Wray, N. R., Visscher, P. M., & Yang, J. (2016). Integration of summary data from GWAS and eQTL studies predicts complex trait gene targets. Nature Genetics, 48(5), 481–487.

