## Supplementary Tables and Figures for "MRLocus: identifying causal genes mediating a trait through Bayesian estimation of allelic heterogeneity"

August 17, 2020

#### Supplementary Tables

| Tissue | eGene | Phenotype | Chr | eSNP | eQTL (bp) | GWAS SNP | GWAS (bp) | $R^2$ |
|---|---|---|---|---|---|---|---|---|
| Artery tibial | MRAS | CAD | 3 | rs13324341 | 138070901 | rs1199338 | 138087467 | 0.983347 |
| Artery tibial | PHACTR1 | CAD | 6 | rs12202891 | 12768218 | rs12202891 | 12768218 | 1 |
| Artery tibial | PHACTR1 | CAD | 6 | rs4711858 | 12880131 | rs4711858 | 12880131 | 1 |
| Artery tibial | PHACTR1 | CAD | 6 | rs9349379 | 12903957 | rs9349379 | 12903957 | 1 |
| Artery tibial | PHACTR1 | CAD | 6 | rs11756003 | 12957621 | rs36049381 | 12953384 | 0.992156 |
| Liver | CETP | HDL | 16 | rs183130 | 56991363 | rs12446515 | 56987015 | 0.985452 |
| Liver | CETP | HDL | 16 | rs289717 | 57009388 | rs4369653 | 56997551 | 0.452439 |
| Liver | CETP | HDL | 16 | rs13337445 | 57026396 | rs17369163 | 57020327 | 0.914334 |
| Liver | LIPC | HDL | 15 | rs13329672 | 58699937 | rs12708454 | 58692202 | 0.46668 |
| Liver | LIPC | HDL | 15 | rs572410 | 58741384 | rs2070895 | 58723939 | 0.531565 |
| Liver | SORT1 | LDL | 1 | rs621414 | 109725239 | rs621414 | 109725239 | 1 |
| Liver | SORT1 | LDL | 1 | rs648673 | 109727284 | rs1337247 | 109703023 | 0.475691 |
| Liver | SORT1 | LDL | 1 | rs11102964 | 109783261 | rs61799430 | 109784082 | 0.420809 |
| Liver | SORT1 | LDL | 1 | rs11102965 | 109784938 | rs1337248 | 109806442 | 0.424259 |
| Liver | SORT1 | LDL | 1 | rs585362 | 109789795 | rs585362 | 109789795 | 1 |
| Liver | SORT1 | LDL | 1 | rs596773 | 109796757 | rs653635 | 109806313 | 0.429544 |
| Liver | SORT1 | LDL | 1 | rs17035665 | 109813719 | rs17035665 | 109813719 | 1 |
| Liver | SORT1 | LDL | 1 | rs4970835 | 109821588 | rs4970835 | 109821588 | 1 |
| Liver | SORT1 | LDL | 1 | rs599839 | 109822166 | rs12740374 | 109817590 | 0.940335 |
| Liver | SORT1 | LDL | 1 | rs17584208 | 109833187 | rs61799430 | 109784082 | 0.495234 |
| Liver | SORT1 | LDL | 1 | rs2272272 | 110009802 | rs12127701 | 109838264 | 0.574156 |
| Liver | SORT1 | LDL | 1 | rs7538453 | 110056537 | rs7538453 | 110056537 | 1 |

Supplementary Table 1: Details on colocalization candidates from eQTL and GWAS summary data employed in MRLocus real data evaluation (see Methods). In addition, all eQTL independent clusters were included in analysis (see LD diagrams in Supplementary Figures 28-32). Genomic locations are in GRCh37 coordinates.

| eQTL Tissue | Resource |
|---|---|
| Artery Tibial (GTEx v8) | https://console.cloud.google.com/storage/browser/gtex-resources |
| Liver | https://www.nature.com/articles/s41598-018-24219-z |

| GWAS Phenotype | Resource |
|---|---|
| CAD (CARDIoGRAMplusC4D) | http://www.cardiogramplusc4d.org/media/cardiogramplusc4d-consortium/data-downloads/cad.additive.Oct2015.pub.zip |
| HDL (UKBB) | https://www.dropbox.com/s/65jisgxwbbdrkaw/30760_irnt.gwas.imputed_v3.both_sexes.tsv.bgz |
| LDL (UKBB) | https://www.dropbox.com/s/4rnjzczwjgs5pgl/30780_irnt.gwas.imputed_v3.both_sexes.tsv.bgz |

Supplementary Table 2: Links to URLs for eQTL and GWAS summary data employed in MRLocus real data evaluation.

### Supplementary Figures

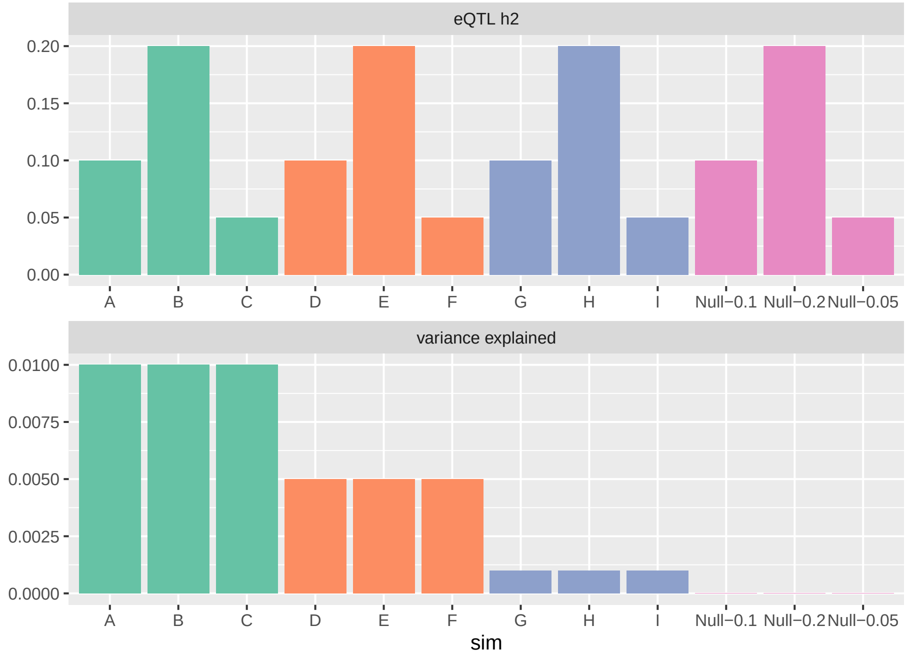


Supplementary Figure 1: Diagram of the 12 types of `twas_sim` simulations performed, varying eQTL $h^2$ (top row) and trait variance explained through gene expression (bottom row), with 20 replicates of each setting resulting in 240 simulations total. Results for simulation A are presented in Figure 2 in the main text, while results for the remaining 11 simulation settings follow in Supplementary Figures.

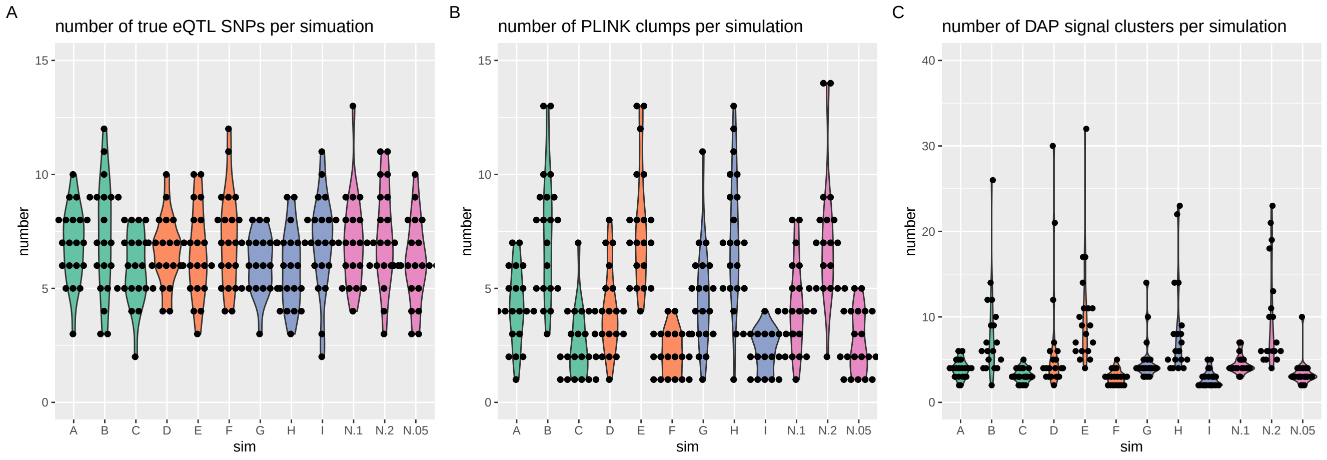


Supplementary Figure 2: Number of (A) true causal eQTL SNPs, (B) PLINK clumps, and (C) DAP signal clusters per simulation.

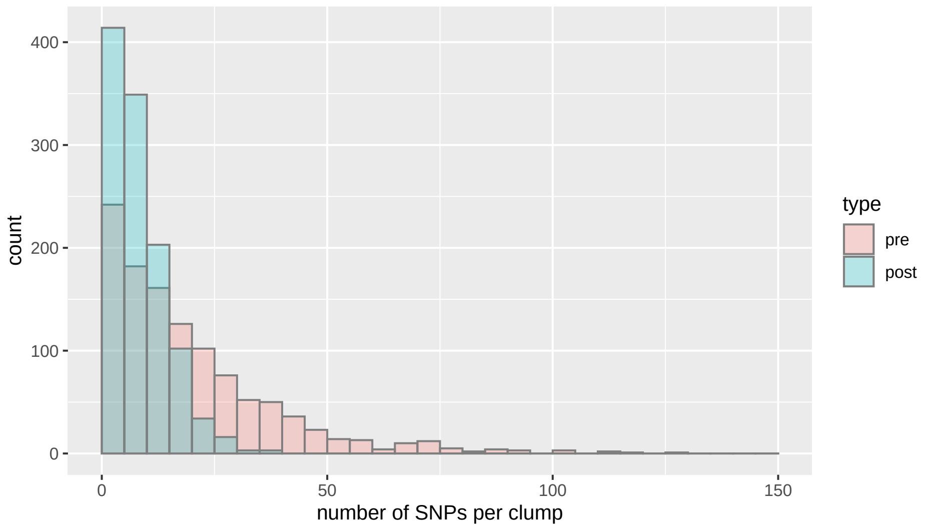


Supplementary Figure 3: SNPs per clump in the 240 simulations, before and after collapsing with MRLocus `collapseHighCorSNPs` function. The mean number of SNPs per clump was 19.9 and 8.8, and the median number of SNPs per clump was 15 and 8 (before and after, respectively).

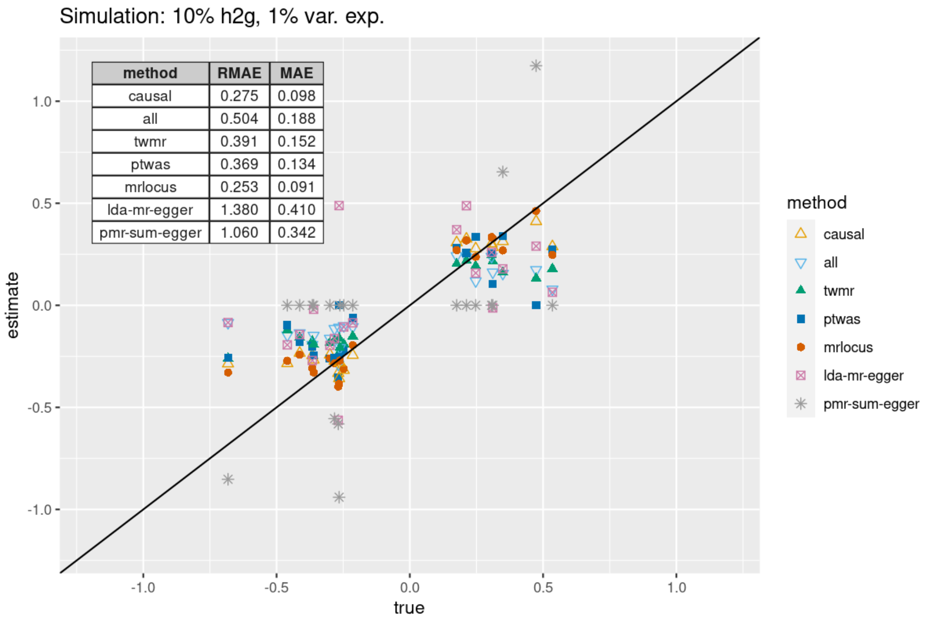


Supplementary Figure 4: Accuracy of gene-to-trait effect estimation in simulation A, including LDA-MR-Egger and PMR-Summary-Egger in comparisons. LDA-MR-Egger did not estimate SE for 6 of the 20 simulations, and PMR-Summary-Egger did not provide an effect estimate for 14 of the 20 simulations.

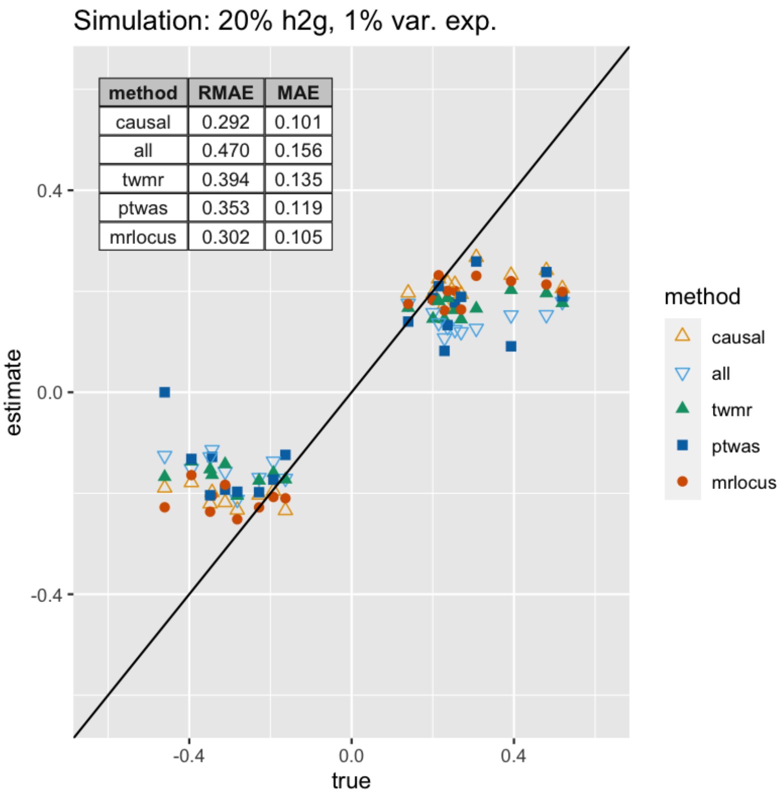


Supplementary Figure 5: Accuracy of gene-to-trait effect estimation in simulation B.

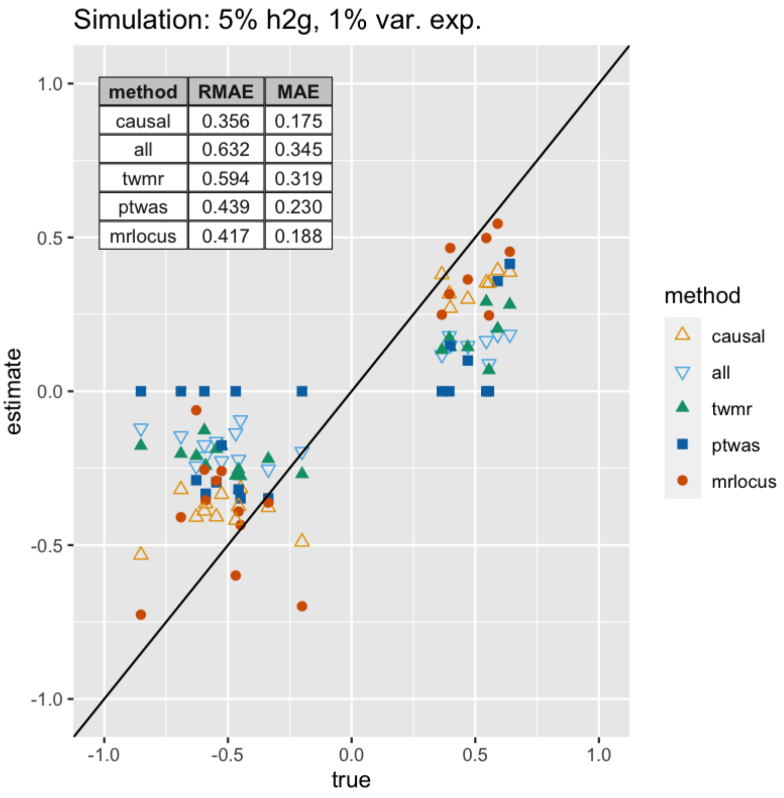


Supplementary Figure 6: Accuracy of gene-to-trait effect estimation in simulation C.

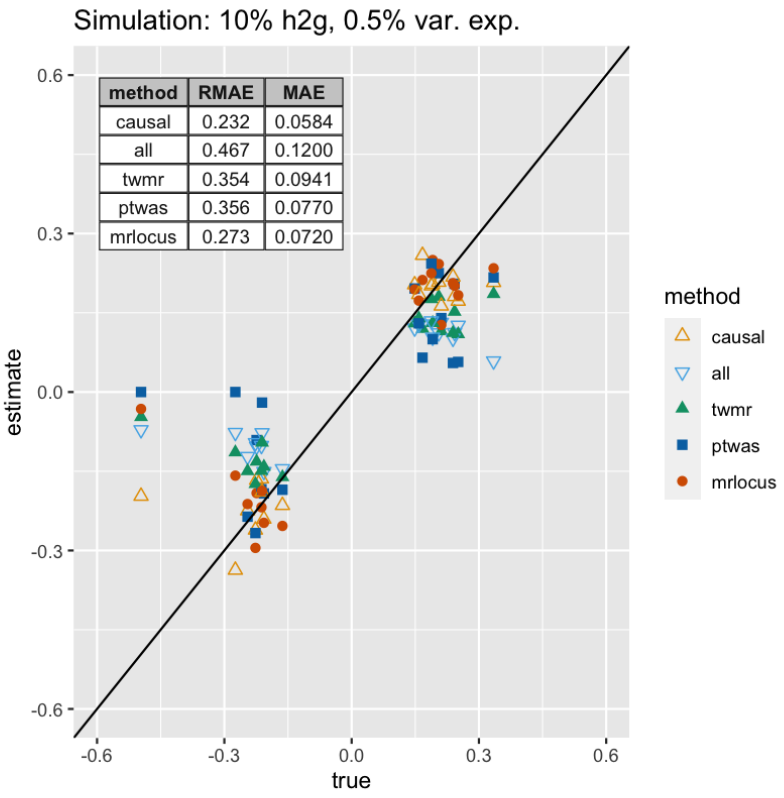


Supplementary Figure 7: Accuracy of gene-to-trait effect estimation in simulation D.

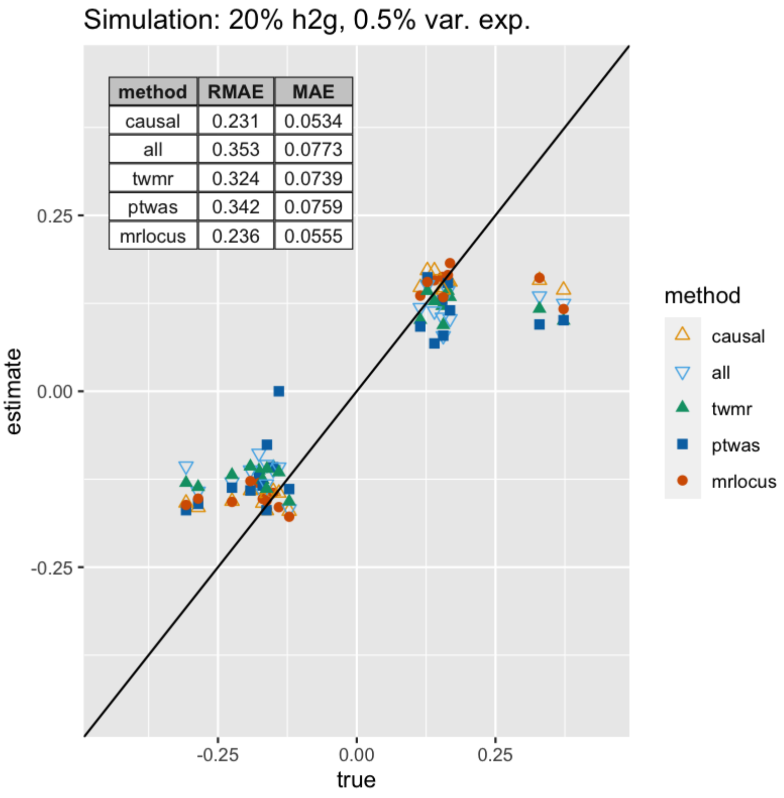


Supplementary Figure 8: Accuracy of gene-to-trait effect estimation in simulation E.

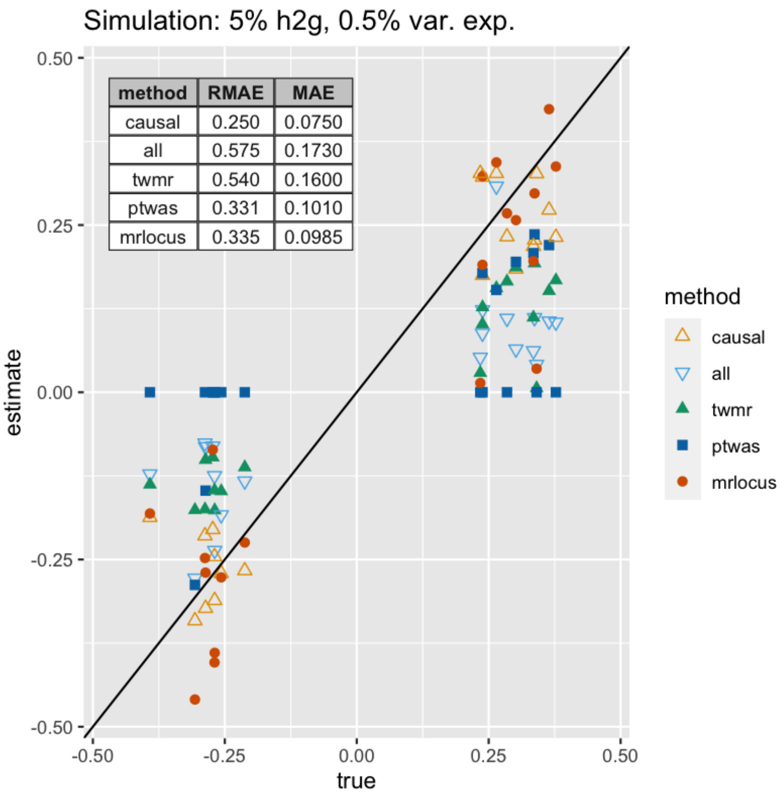


Supplementary Figure 9: Accuracy of gene-to-trait effect estimation in simulation F.

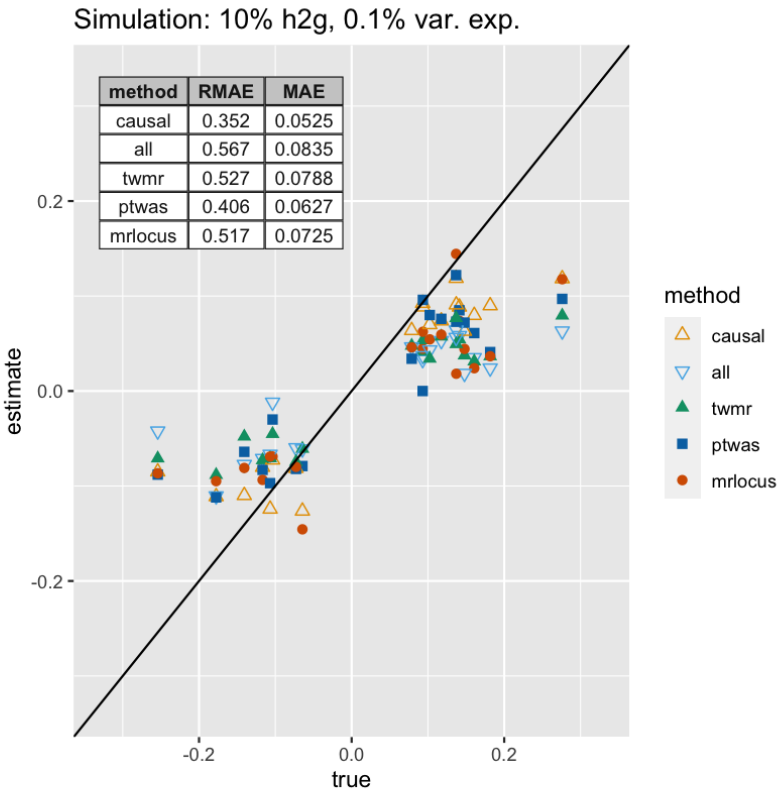


Supplementary Figure 10: Accuracy of gene-to-trait effect estimation in simulation G.

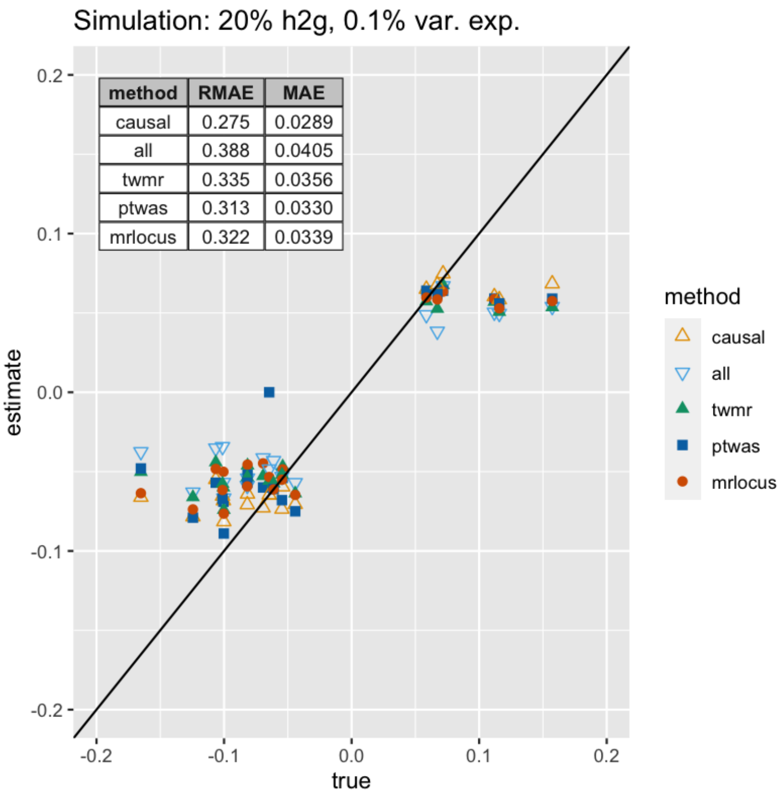


Supplementary Figure 11: Accuracy of gene-to-trait effect estimation in simulation H.

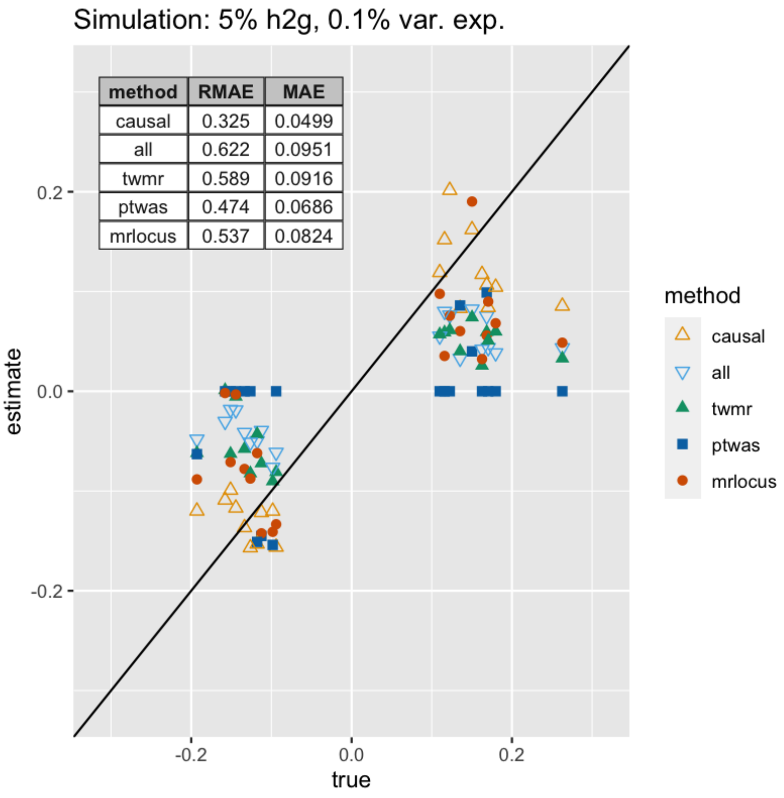


Supplementary Figure 12: Accuracy of gene-to-trait effect estimation in simulation I.

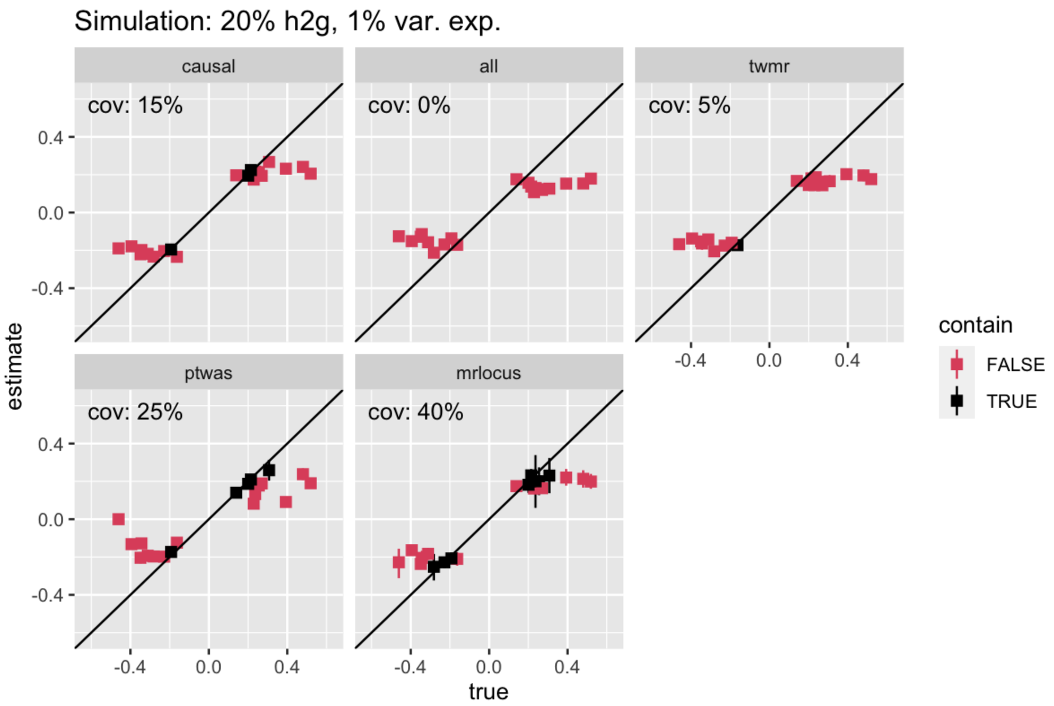


Supplementary Figure 13: Coverage of confidence or credible intervals for the gene-to-trait effect in simulation B.

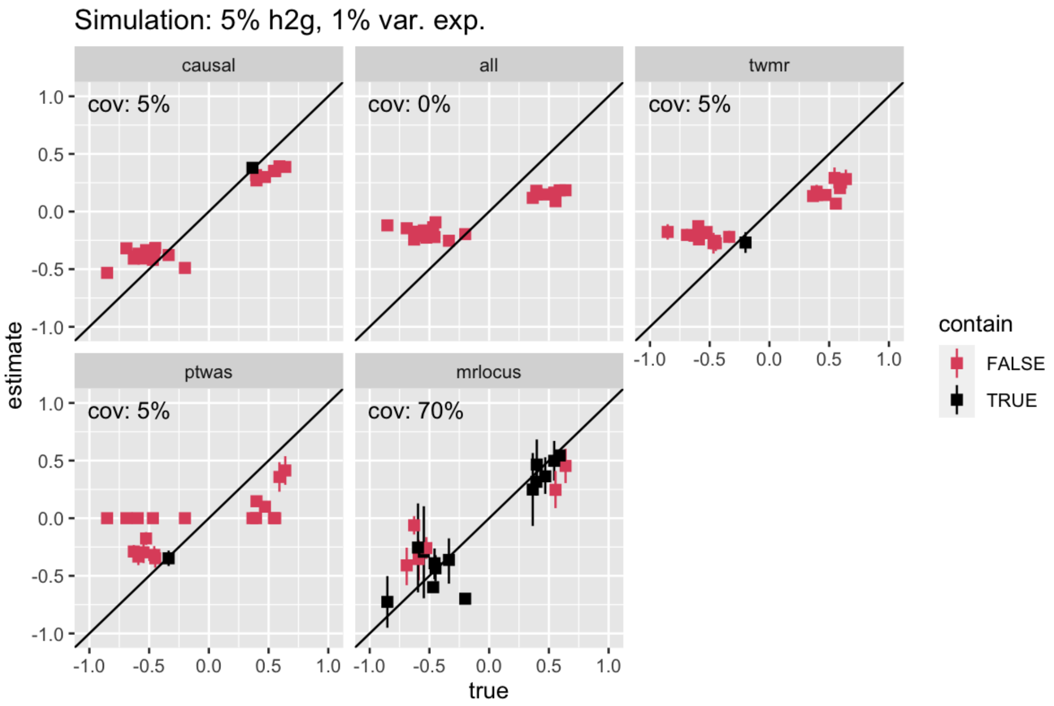


Supplementary Figure 14: Coverage of confidence or credible intervals for the gene-to-trait effect in simulation C.

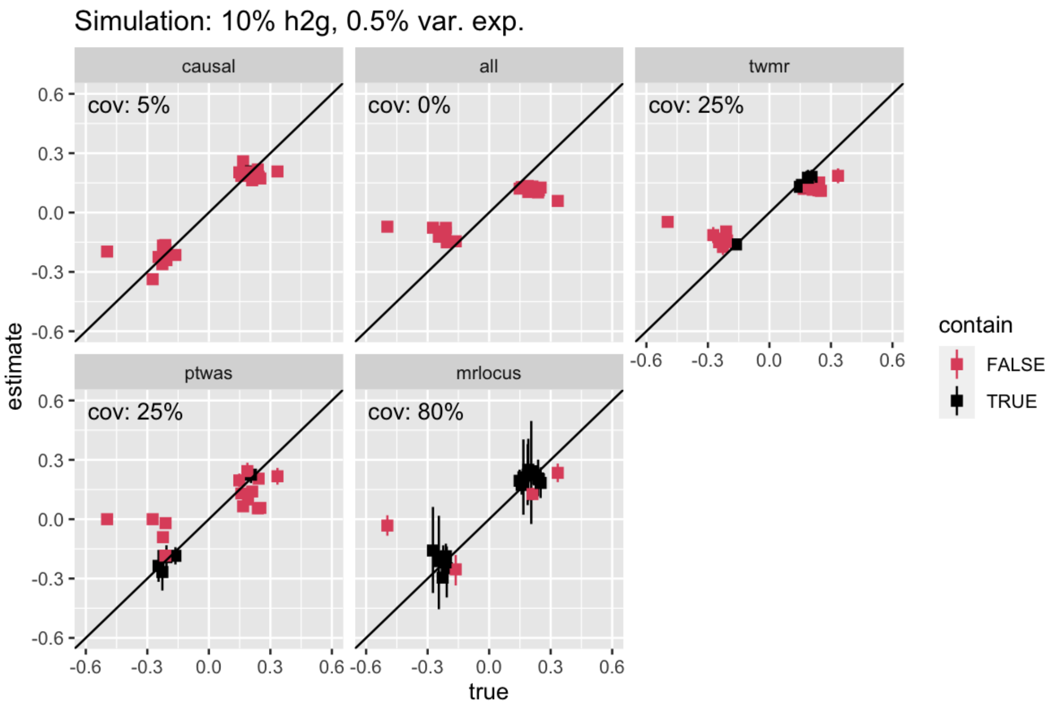


Supplementary Figure 15: Coverage of confidence or credible intervals for the gene-to-trait effect in simulation D.

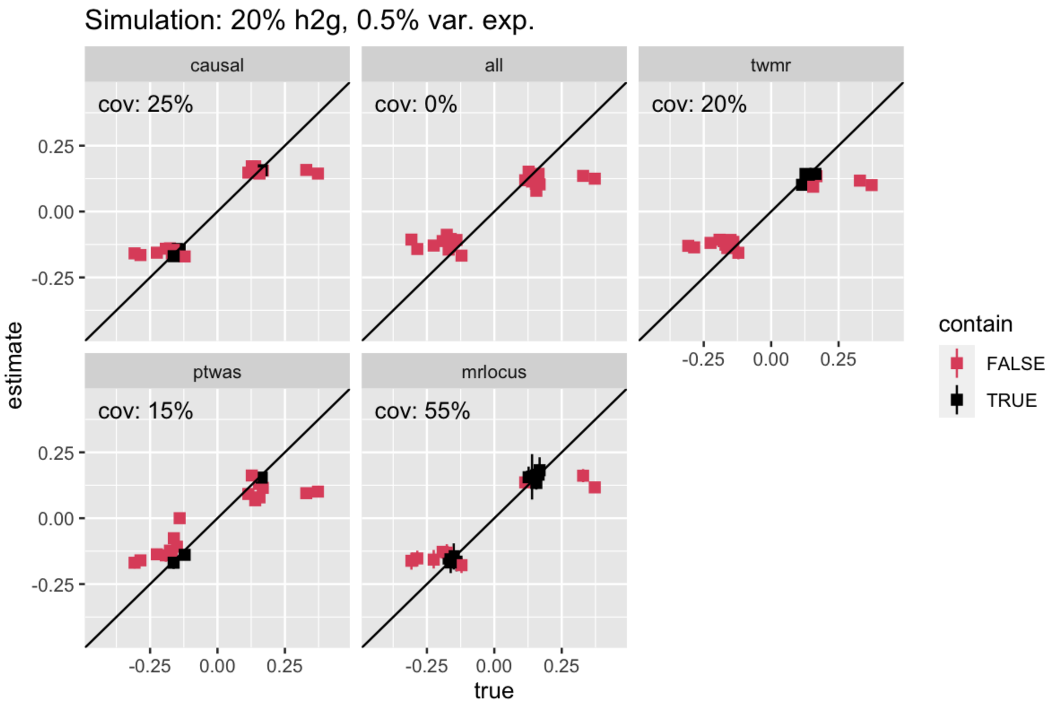


Supplementary Figure 16: Coverage of confidence or credible intervals for the gene-to-trait effect in simulation E.

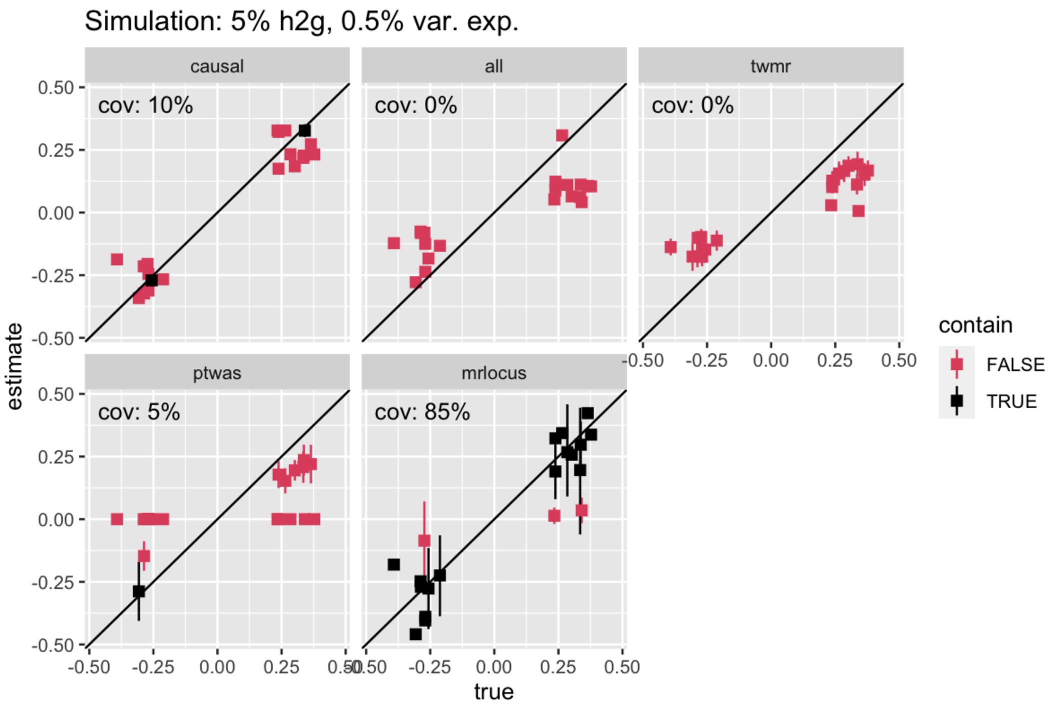


Supplementary Figure 17: Coverage of confidence or credible intervals for the gene-to-trait effect in simulation F.

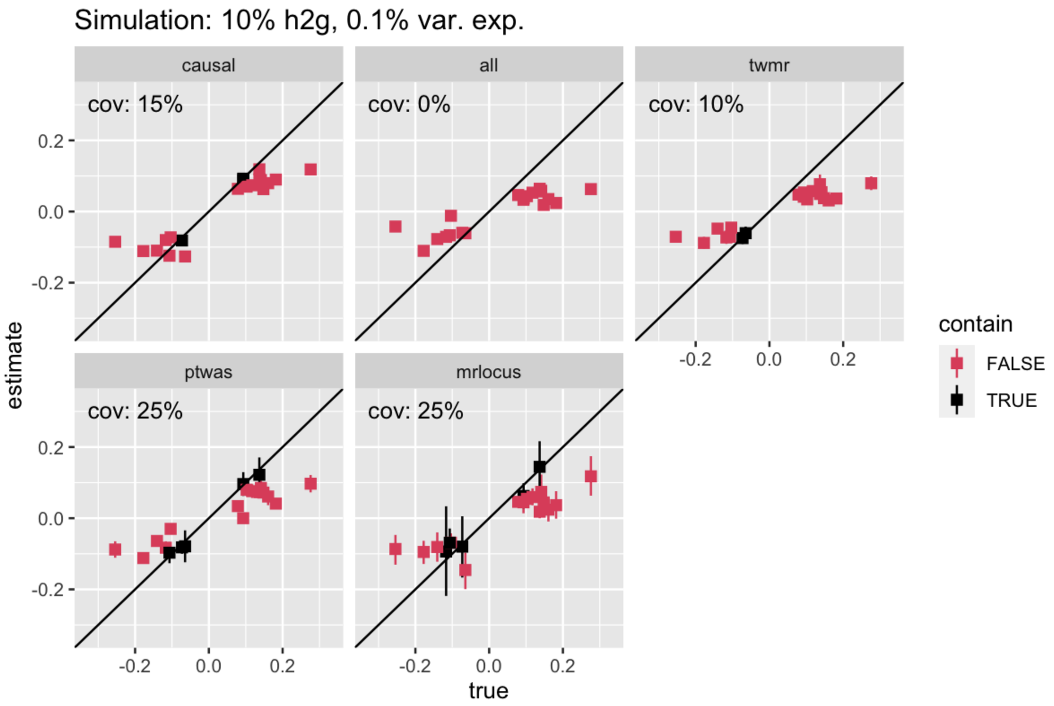


Supplementary Figure 18: Coverage of confidence or credible intervals for the gene-to-trait effect in simulation G.

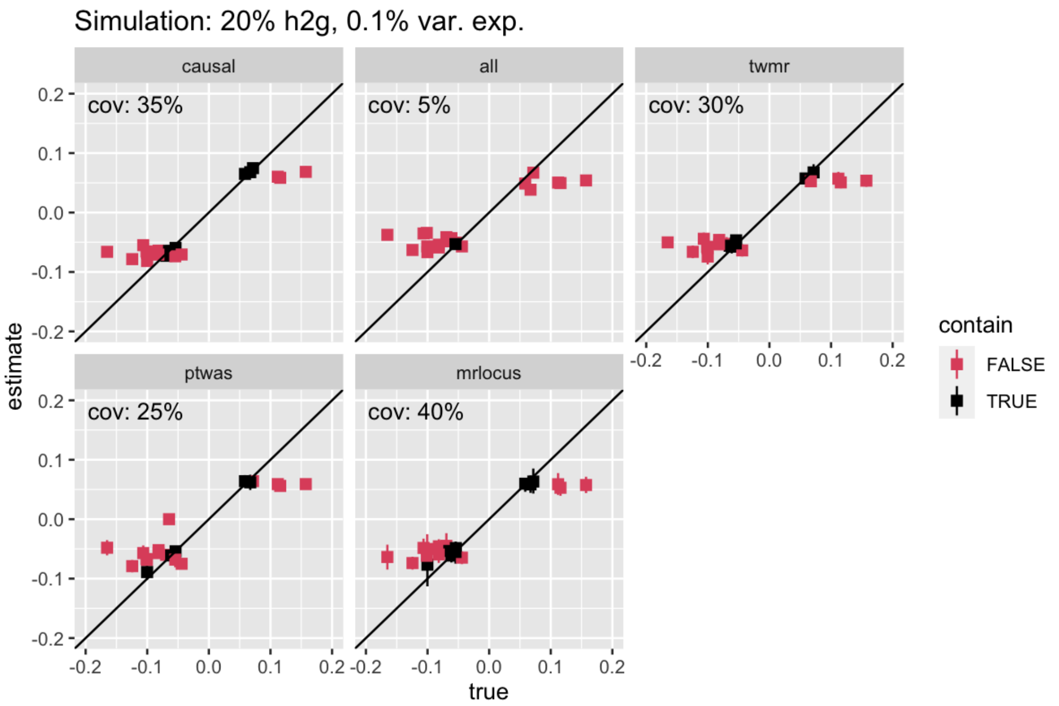


Supplementary Figure 19: Coverage of confidence or credible intervals for the gene-to-trait effect in simulation H.

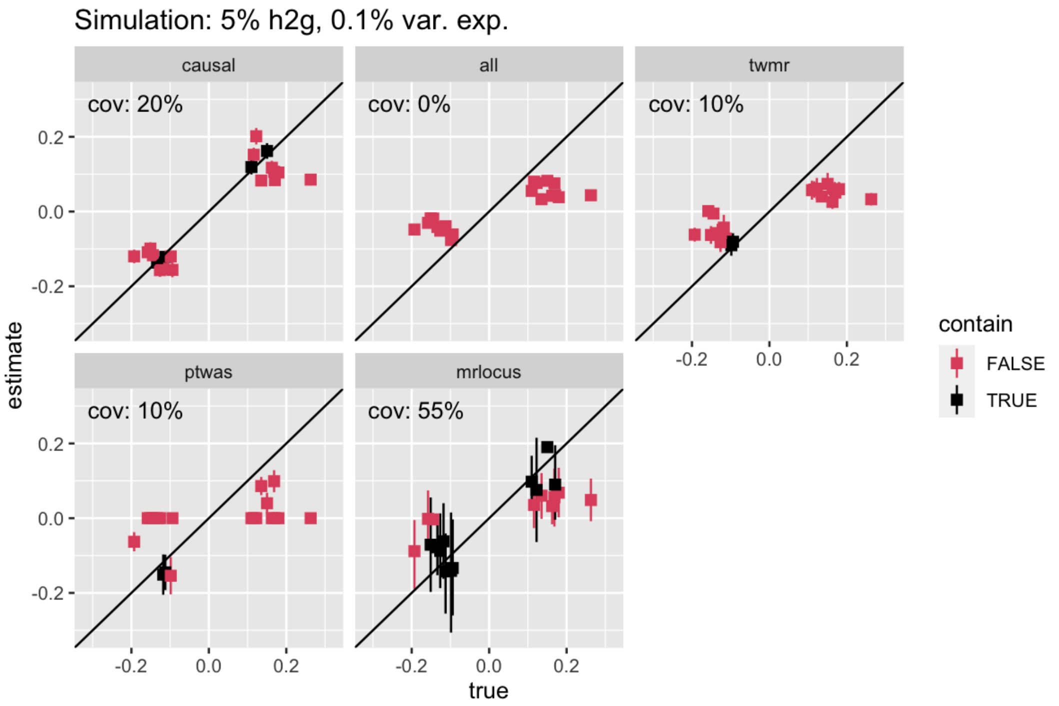


Supplementary Figure 20: Coverage of confidence or credible intervals for the gene-to-trait effect in simulation I.

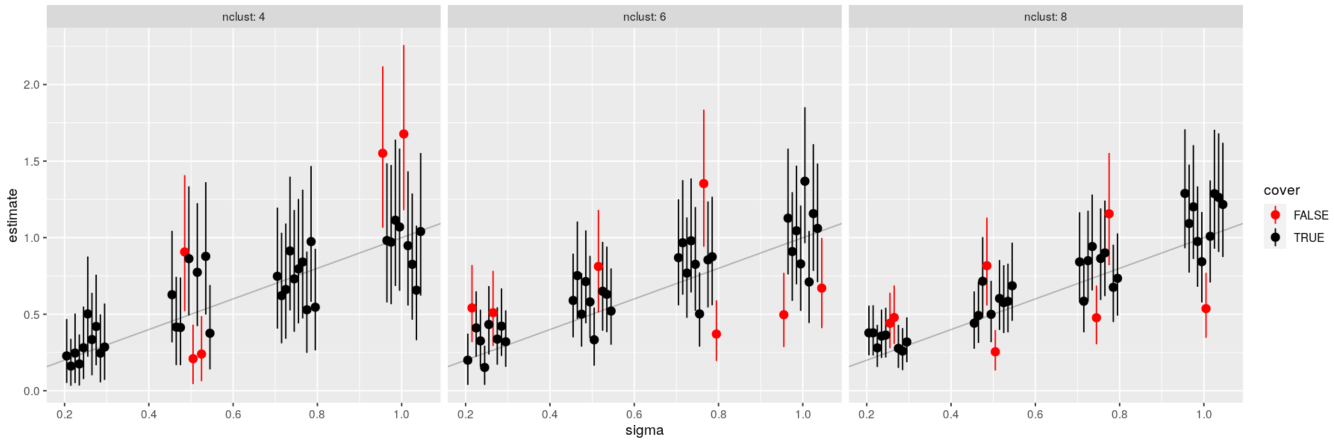


Supplementary Figure 21: Simulation assessing MRLocus' estimation of the scale of dispersion ($\sigma$) across independent signal clusters. The summary statistics for eQTL and GWAS were generated from a multivariate normal distribution as in the eCAVIAR model, using a simulated LD matrix. The true slope ($\alpha$) was set to 1, the true $\sigma$ varied from 0.25 to 1 in increments of 0.25 (x-axis), and the number of LD-independent clusters varied between 4, 6, and 8 (left, middle, and right panels), with 10 iterations per setting (plotted with spacing to avoid overplotting). The posterior mean is indicated with a dot, while 80% quantile-based credible intervals and their coverage of the true value are indicated with the line and its color. The simulation script is included within the MRLocus package test directory, as an R script `"test_sigma.R"`.

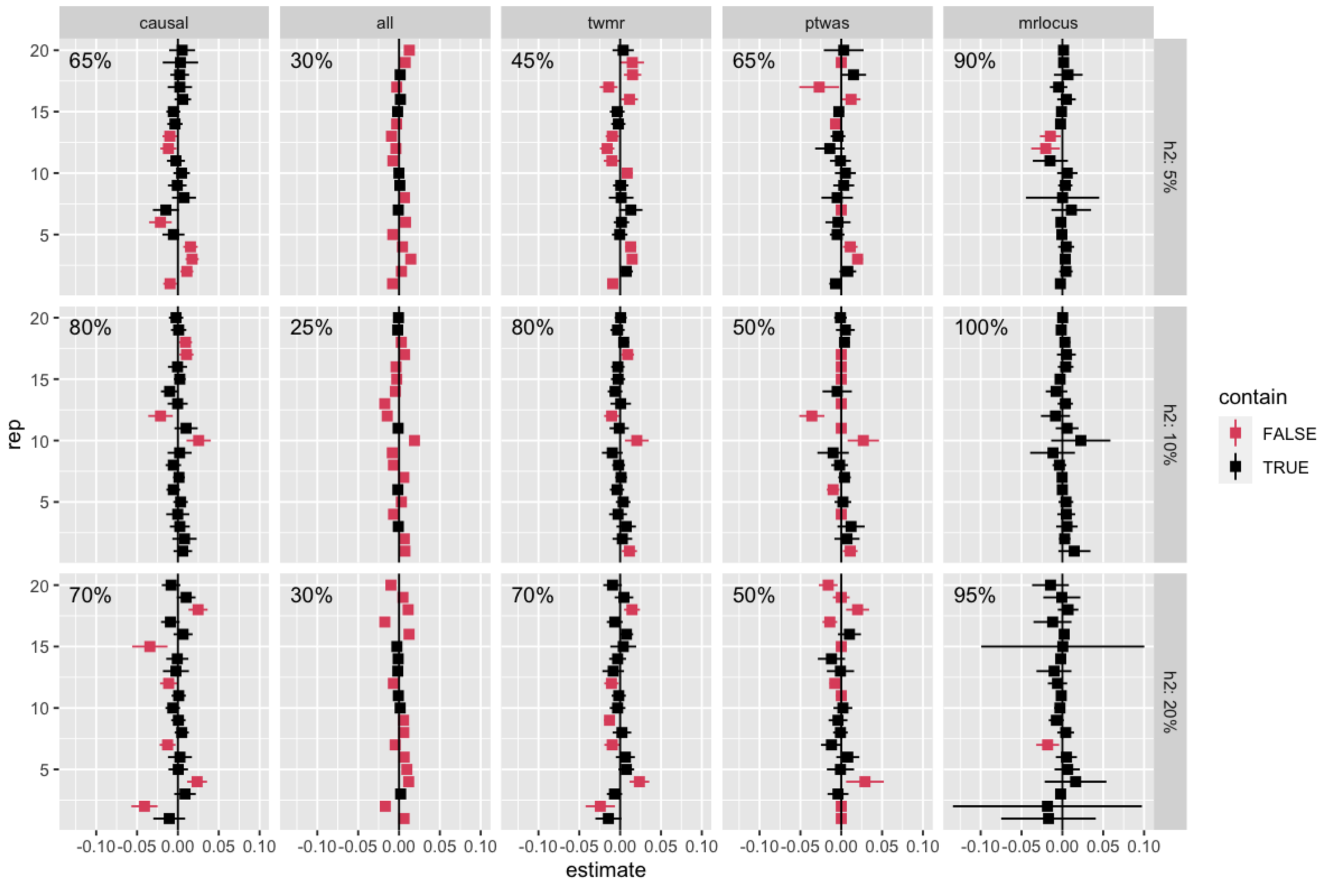


Supplementary Figure 22: Coverage of confidence or credible intervals for the 3 null simulation settings.

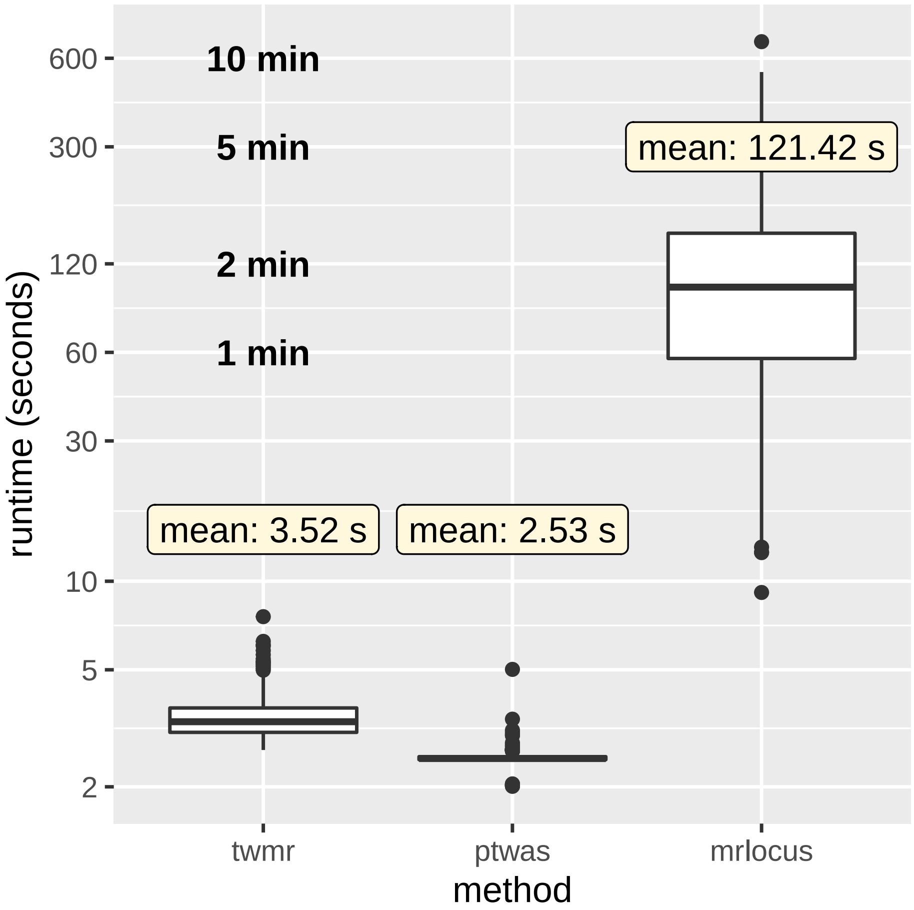


Supplementary Figure 23: Runtime for TWMR, PTWAS, and MRLocus on the 240 simulations. The runtime for a single locus using a single core is shown on the y-axis (log scale).

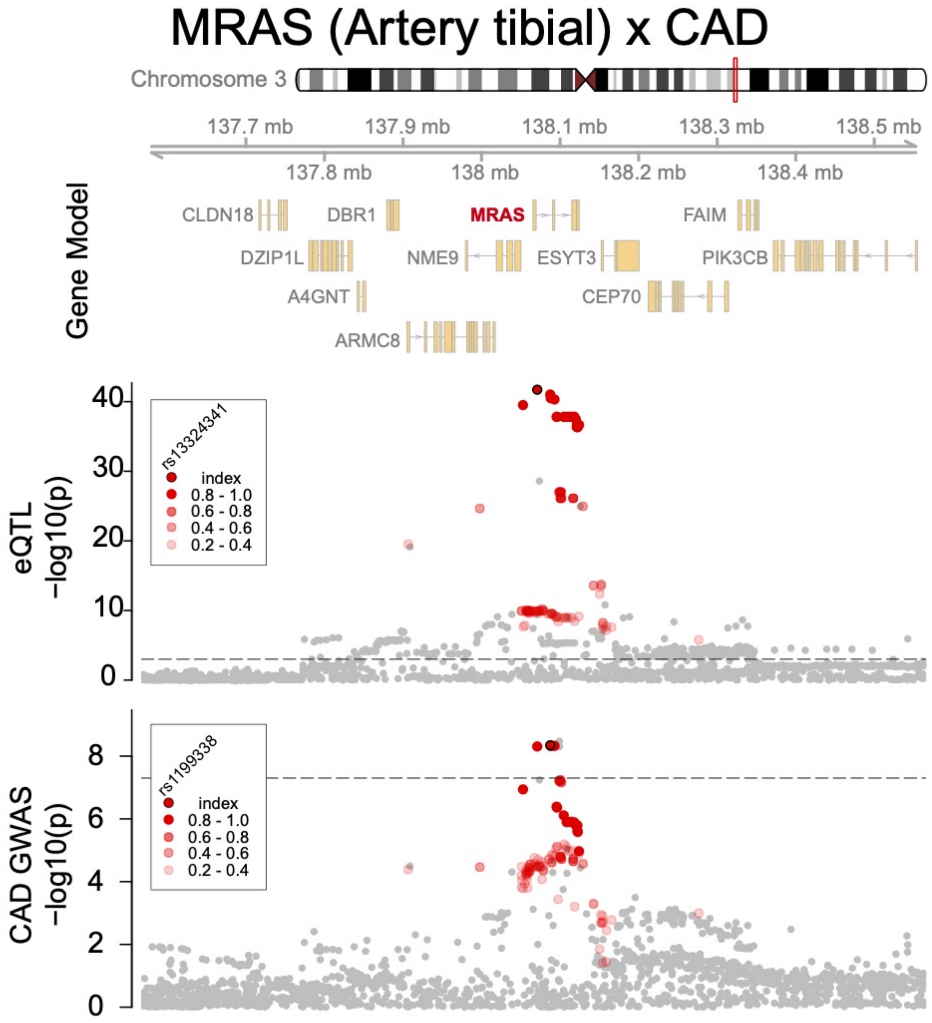


Supplementary Figure 24: Colocalized signals in the *MRAS* region. From top panel to bottom, gene model (NCBI Refseq), eQTL for MRAS in artery tibial (GTEx; N = 663) and CAD association within CARDIoGRAMplusC4D ($N_{\text{case}}$ = 60,801 and $N_{\text{control}}$ = 123,504) (M. Nikpay et al., 2015). LD was calculated to independent SNPs within 1KG EUR and colored accordingly. Symbols indicate independent co-localized ($r^2 > 0.4$) eQTL-GWAS pairs. Dashed line indicates a significance threshold at p = 0.001 or p = 5x$10^{-8}$ for eQTL and GWAS respectively.

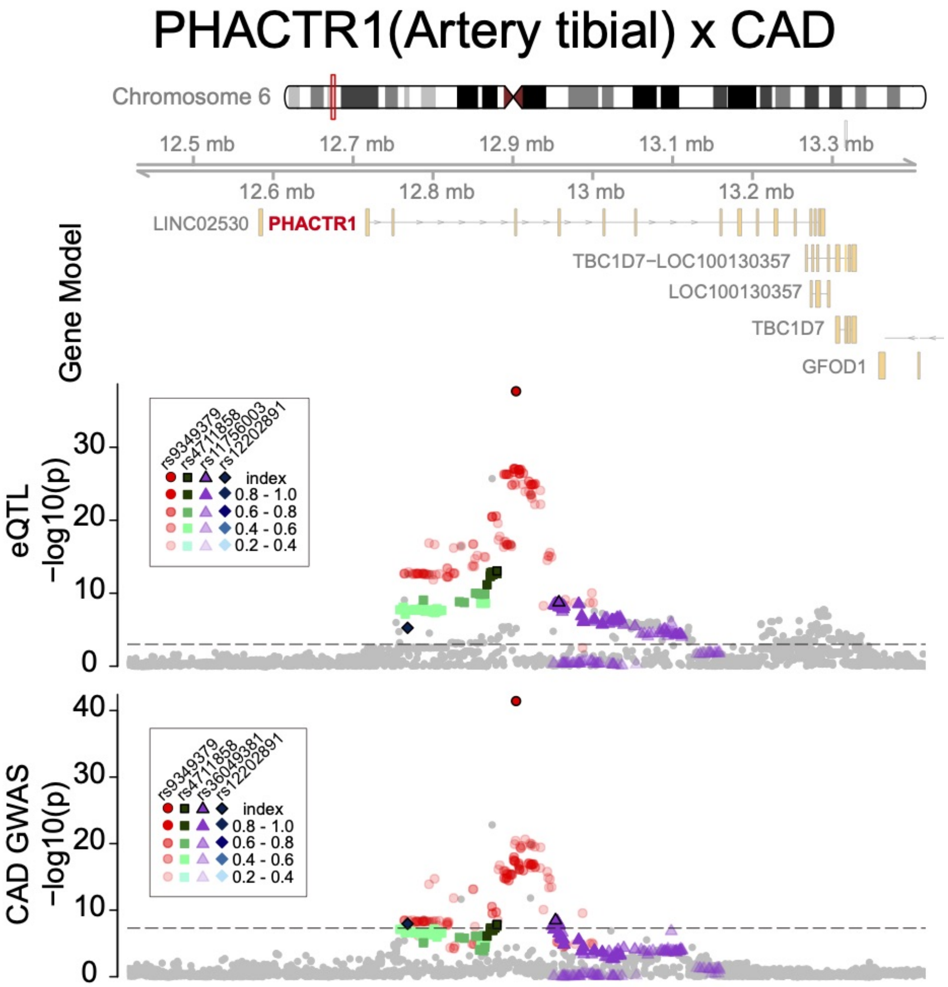


Supplementary Figure 25: Colocalized signals in the *PHACTR1* region. From top panel to bottom, gene model (NCBI Refseq), eQTL for PHACTR1 in artery tibial (GTEx; N = 663) and CAD association within CARDIoGRAMplusC4D ($N_{\text{case}}$ = 60,801 and $N_{\text{control}}$ = 123,504) (M. Nikpay et al., 2015). LD was calculated to independent SNPs within 1KG EUR and colored accordingly. Symbols indicate independent co-localized ($r^2 > 0.4$) eQTL-GWAS pairs. Dashed line indicates a significance threshold at p = 0.001 or p = 5x$10^{-8}$ for eQTL and GWAS respectively.

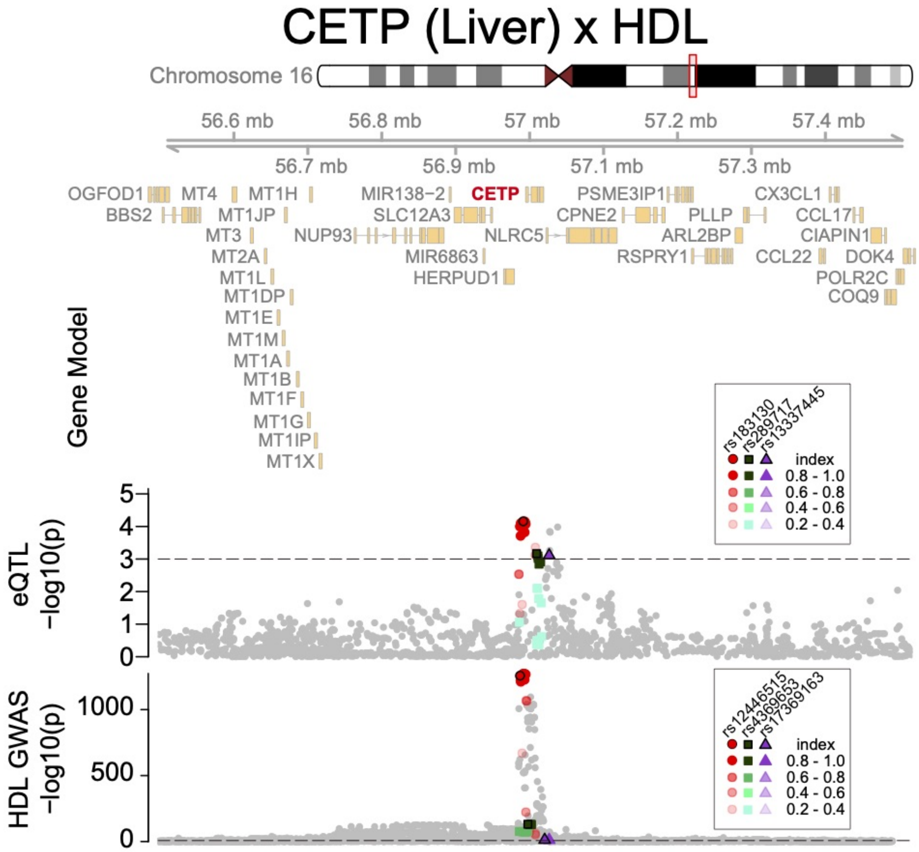


Supplementary Figure 26: Colocalized signals in the *CETP* region. From top panel to bottom, gene model (NCBI Refseq), eQTL for CETP in liver (N = 588) (Strunz et al., 2018) and HDL association within UKBB (N = 315,133). LD was calculated to independent SNPs within 1KG EUR and colored accordingly. Symbols indicate independent co-localized ($r^2 > 0.4$) eQTL-GWAS pairs. Dashed line indicates a significance threshold at p = 0.001 or p = $5 \times 10^{-8}$ for eQTL and GWAS respectively.

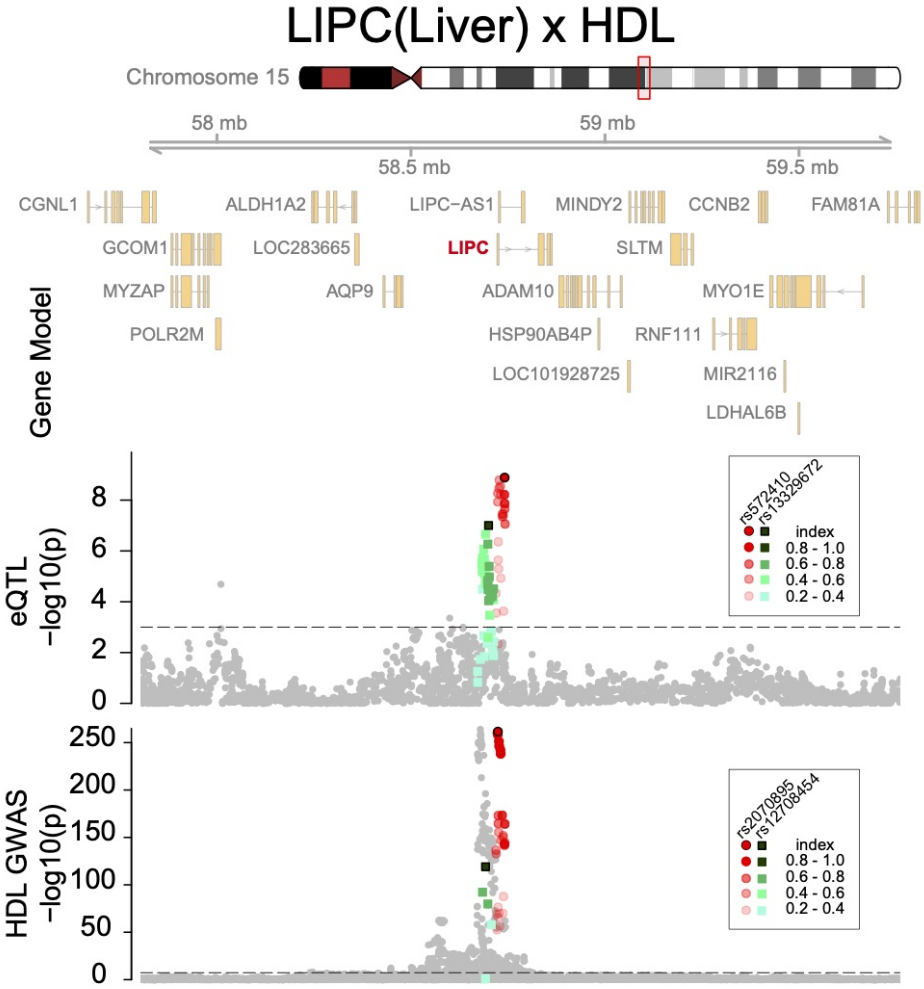


Supplementary Figure 27: Colocalized signals in the *LIPC* region. From top panel to bottom, gene model (NCBI Refseq), eQTL for LIPC in liver (N = 588) (Strunz et al., 2018) and HDL association within UKBB (N = 315,133). LD was calculated to independent SNPs within 1KG EUR and colored accordingly. Symbols indicate independent co-localized ($r^2 > 0.4$) eQTL-GWAS pairs. Dashed line indicates a significance threshold at p = 0.001 or p = $5x10^{-8}$ for eQTL and GWAS respectively.

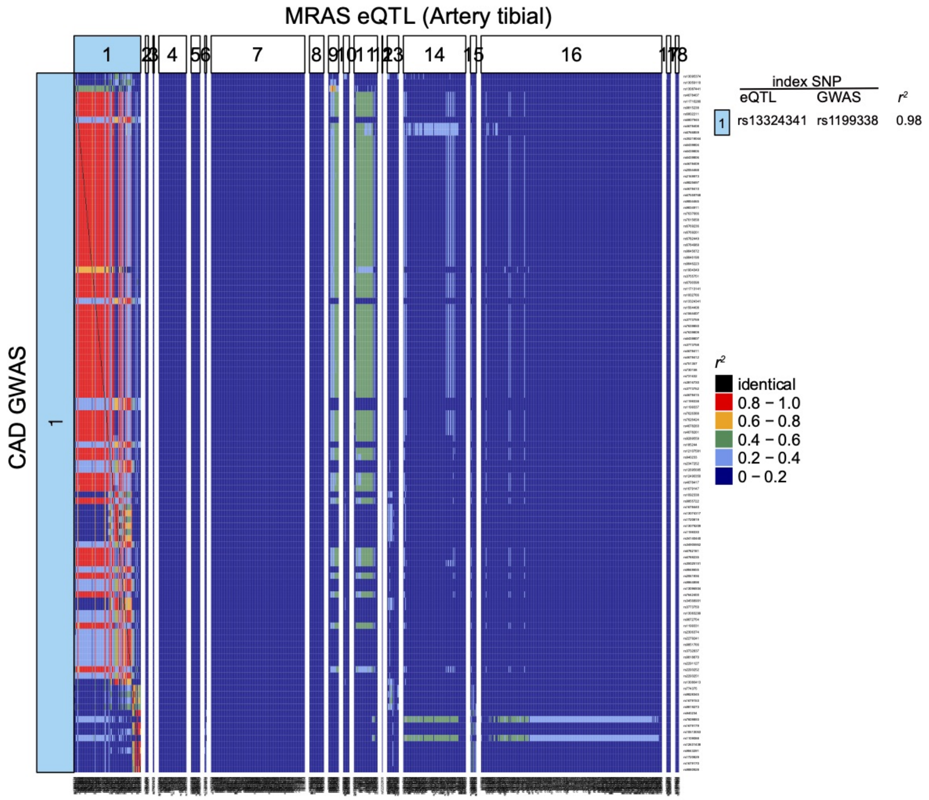


Supplementary Figure 28: LD pattern across independent clusters from *MRAS* eQTL (Artery tibial) and CAD GWAS. LD ($r^2$) between SNPs in independent clumps from *MRAS* in artery tibial (columns) and CAD GWAS SNPs (rows). Color bars at the top and left represent cluster (pair) ID which is sorted by base position of index eQTL SNPs. LD was calculated to independent SNPs within 1KG EUR and colored accordingly.

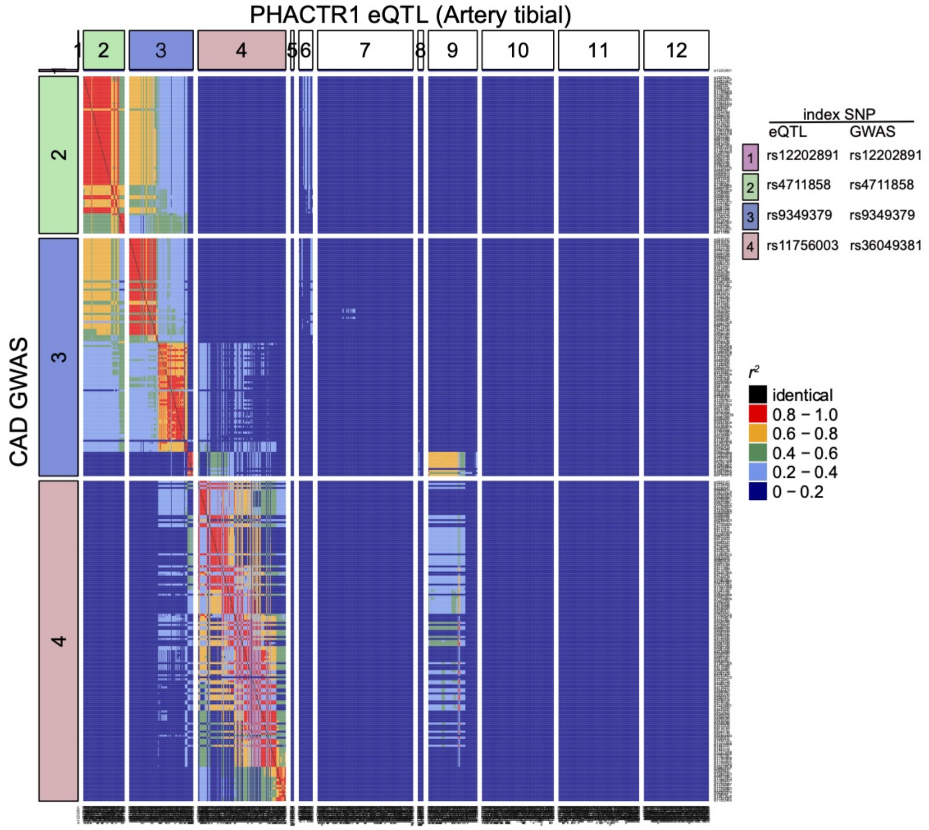


Supplementary Figure 29: LD pattern across independent clusters from *PHACTR1* eQTL (Artery tibial) and CAD GWAS. LD ($r^2$) between SNPs in independent clumps from *PHACTR1* in artery tibial (columns) and CAD GWAS SNPs (rows). Color bars at the top and left represent cluster (pair) ID which is sorted by base position of index eQTL SNPs. LD was calculated to independent SNPs within 1KG EUR and colored accordingly.

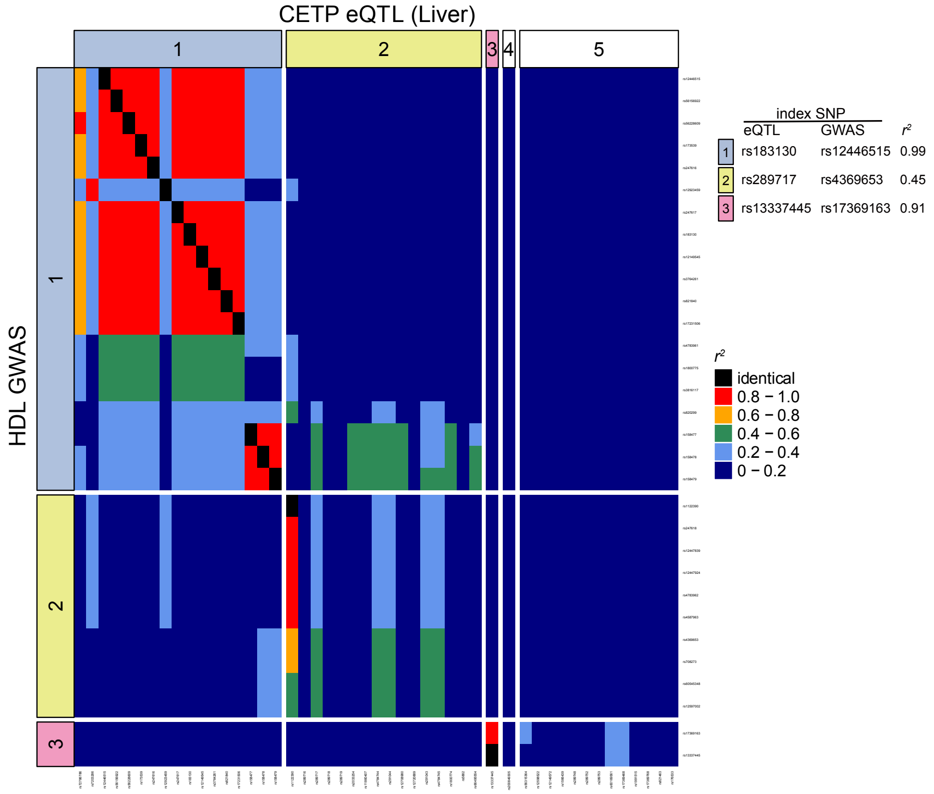


Supplementary Figure 30: LD pattern across independent clusters from *CETP* eQTL (liver) and HDL GWAS. LD ($r^2$) between SNPs in independent clumps from *CETP* in liver (columns) and HDL GWAS SNPs (rows). Color bars at the top and left represent cluster (pair) ID which is sorted by base position of index eQTL SNPs. LD was calculated to independent SNPs within 1KG EUR and colored accordingly.

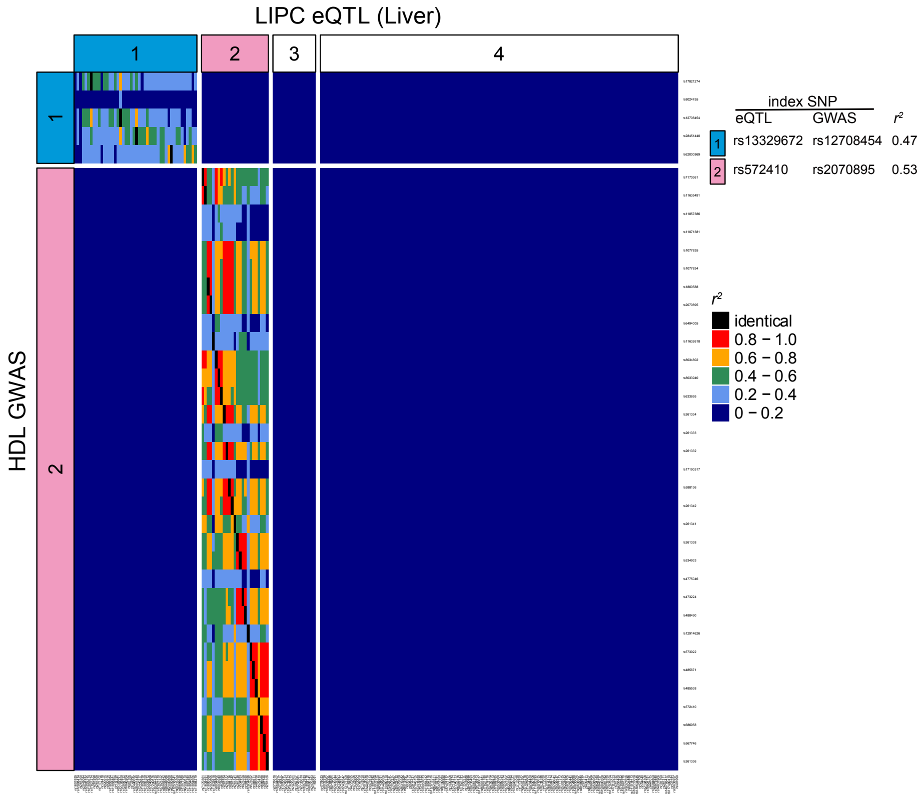


Supplementary Figure 31: LD pattern across independent clusters from *LIPC* eQTL (liver) and HDL GWAS. LD ($r^2$) between SNPs in independent clumps from *LIPC* in liver (columns) and HDL GWAS SNPs (rows). Color bars at the top and left represent cluster (pair) ID which is sorted by base position of index eQTL SNPs. LD was calculated to independent SNPs within 1KG EUR and colored accordingly.

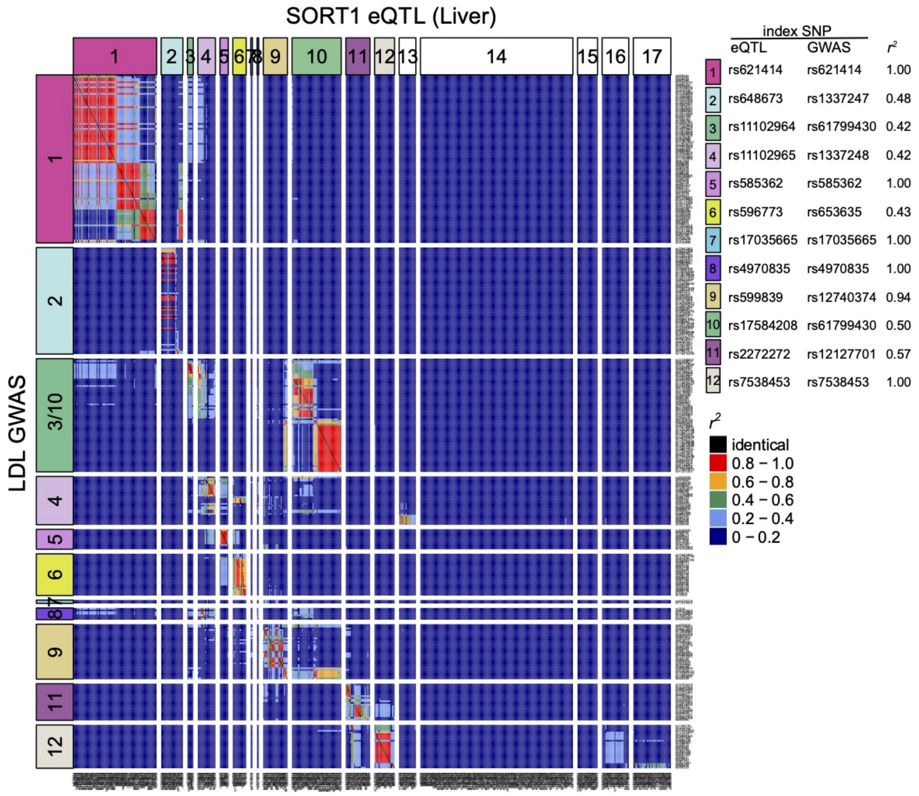


Supplementary Figure 32: LD pattern across independent clusters from *SORT1* eQTL (liver) and LDL GWAS. LD ($r^2$) between SNPs in independent clumps from *SORT1* in liver (columns) and LDL GWAS SNPs (rows). Color bars at the top and left represent cluster (pair) ID which is sorted by base position of index eQTL SNPs. LD was calculated to independent SNPs within 1KG EUR and colored accordingly.

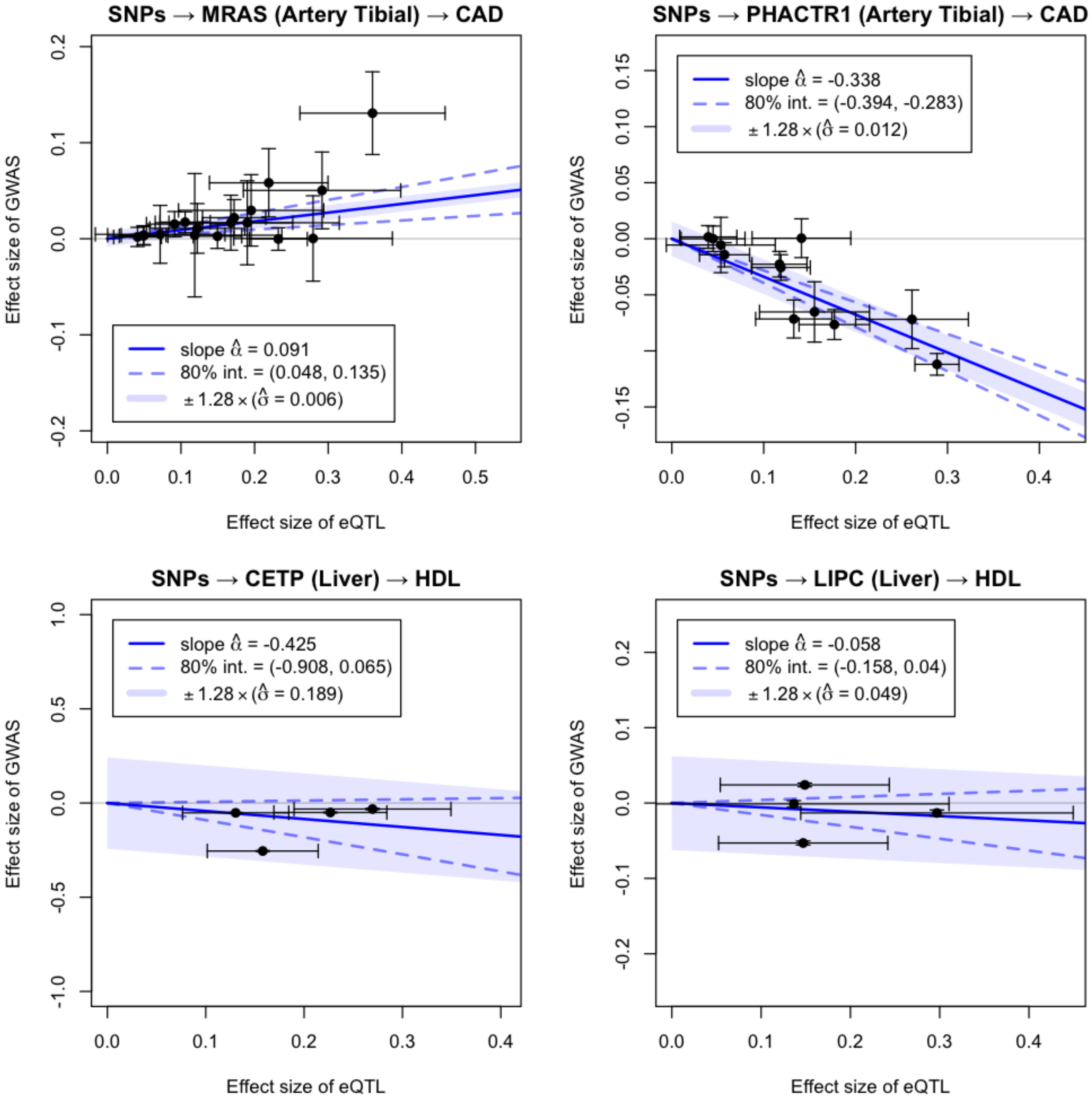


Supplementary Figure 33: MRLocus plots for the four additional gene-trait pairs, in addition to *SORT1* in Figure 4.
