## Supplementary Methods for "MRLocus: identifying causal genes mediating a trait through Bayesian estimation of allelic heterogeneity"

August 13, 2020

#### 1 MRLocus statistical model

MRLocus proceeds in two separate hierarchical models, which are encoded in the Stan programming language and with posterior inference performed using the Stan and RStan software packages [Carpenter et al., 2017, Stan Development Team, 2020]. In section 1.1 we define the model for the colocalization step, and in section 1.2 we define the model for the slope fitting step.

##### 1.1 Colocalization step

###### 1.1.1 Input data

The first step performs colocalization of eQTL (A) and GWAS (B) signals across a number of LD-independent signal clusters $j \in 1, \dots, J$, using summary statistics from both studies: $\widehat{\beta}^A_{i,j}$ and $\text{se}(\widehat{\beta}^A_{i,j})$, $\widehat{\beta}^B_{i,j}$ and $\text{se}(\widehat{\beta}^B_{i,j})$ for SNP $i \in 1, \dots, n_j$ in cluster $j$ for study A and B, and the respective LD matrices for each cluster $j$: $\Sigma^A_j$ and $\Sigma^B_j$. The clusters and the individual SNPs per cluster must be matched across study, and our procedure for performing this matching is described in the Methods section in the main text. The estimated coefficients $\widehat{\beta}^X_{i,j}$ for $X \in \{A, B\}$ refer to either the estimated coefficients from a linear model of a continuous trait $y$ on genotype dosages $\{0, 1, 2\}$, or the estimated log odds from a logistic regression of a binary trait $y$ modeled on genotype dosages.

While it would be preferable to use allelic fold change (aFC) [Mohammadi et al., 2017] or ACME effect sizes [Palowitch et al., 2018] for the eQTL (A) study in MRLocus modeling, in practice we typically are provided with publicly available estimated coefficients representing inverse normal transformed (INT) or $\log_2$ transformed expression values regressed on genotype dosages. For eQTL coefficients derived from INT expression data, the mediation effect estimated by MRLocus represents the effect on the trait from modifying gene expression by 1 SD, while for eQTL coefficients derived from $\log_2$ transformed expression data, the mediation effect represents the effect on the trait from doubling gene expression.

The eQTL and GWAS studies are referred to as "A" and "B" in the formula and code below for generalization, for example, the eQTL study could be replaced with a pQTL (protein quantitative trait loci) study. "A" therefore refers to a study of a trait (or "exposure") that is believed to be causally upstream of the trait examined in study "B" (or "outcome").

###### 1.1.2 Collapsing and allele flipping

MRLocus contains two convenience functions, `collapseHighCorSNPs` and `flipAllelesAndGather`, which are described briefly. The first function uses hierarchical clustering based on the LD matrix (the user must pick which to use if two are available), in order to collapse SNPs into groups using complete linkage, and thresholding the resulting dendrogram at 0.95 correlation. The SNP with the highest absolute $z$-score within a collapsed set is chosen as the representative SNP.

`flipAllelesAndGather` performs a number of allele flipping steps for assisting statistical modeling and visualization. The alleles are flipped such that the index SNP (as defined by its absolute value of $z$-score) for study A has a positive estimated coefficient. This is to simplify the interpretations of the plots – such that we are always describing the effects on downstream traits for expression increasing alleles. Additionally, we flip alleles such that SNPs with positive correlation of genotypes in either $\Sigma_j^A$ or $\Sigma_j^B$ (the user must pick which to use) are kept the same, while SNPs with negative correlation of genotypes have their alleles flipped. Allele flipping involves both keeping track of the reference and effect allele, as well as changing the sign of the estimated coefficient. This function also performs checks such that the A and B study agree in terms of the effect and reference allele. If two LD matrices are provided, one is prioritized for generating positive correlations of genotypes, while the other has its alleles flipped for consistency.

##### 1.1.3 Scaling

In practice, before supplying $\widehat{\beta}_{i,j}^A$ and $\widehat{\beta}_{i,j}^B$ and the associated standard errors to the colocalization hierarchical model, the values are scaled such that the index SNP (as defined by its absolute value of $z$-score) for study A has estimated coefficient of $\pm 1$ for both study A and B. If two or more SNPs have the same $z$-score, the first is chosen. This simplifies the Stan code and improves model fit, as the two studies are then at comparable scale. The scaling is reversed after the Stan model is fit. For the user-input estimated coefficients and standard errors for study $X \in \{A, B\}$ and cluster $j$, $\widehat{\beta}_{i,j}^{X,input}$ and $\text{se}(\widehat{\beta}_{i,j}^{X,input})$, in the first step we create scaled estimated coefficients:

$$z_j^* = \max_i \left( |\widehat{\beta}_{i,j}^{A,input}| / \text{se}(\widehat{\beta}_{i,j}^{A,input}) \right) \tag{1}$$

$$i_j^* = \min\,(i \in 1, \ldots, n_j) \;\; s.t. \; |\widehat{\beta}_{i,j}^{A,input}| / \text{se}(\widehat{\beta}_{i,j}^{A,input}) = z_j^* \tag{2}$$

$$s_j^X = 1/|\widehat{\beta}_{i^*,j}^{X,input}| \tag{3}$$

$$\widehat{\beta}_{i,j}^X = s_j^X \, \widehat{\beta}_{i,j}^{X,input}, \; i \in 1, \ldots, n_j \tag{4}$$

$$\text{se}(\widehat{\beta}_{i,j}^X) = s_j^X \, \text{se}(\widehat{\beta}_{i,j}^{X,input}), \; i \in 1, \ldots, n_j \tag{5}$$

Note that equations (1-2) refer specifically to study A, while equations (3-5) refer to steps that are repeated for $X = A$ and $X = B$. Again, in words, $i_j^*$ is the index of the first occurrence of the maximal value of absolute value of $z$-score in study A and cluster $j$.

##### 1.1.4 Colocalization

Colocalization refers to the task of determining if the same signal in eQTL and GWAS summary statistics arise from the same causal variant(s), considering the correlation of genotypes in a locus (the LD matrix). Here we perform colocalization using a generative model for the estimated coefficients, where the true coefficients will be modeled and their posterior distribution used for inference. In the following equations, $\widehat{\beta}_{i,j}^X$ and $\text{se}(\widehat{\beta}_{i,j}^X)$ refer to these scaled estimates as described in the previous section, but in the following section 1.2 on slope fitting, the values are transformed back to the original scale by multiplying by $1/s_j^X$ for $X \in \{A, B\}$ respectively.

We use a statistical model for the summary statistics motivated by the eCAVIAR model [Hormozdiari et al., 2016]. In eCAVIAR, the summary statistic $z$-scores in a vector $S$ are modeled as a multivariate normal distribution with mean vector $\Sigma\Lambda$ and covariance matrix $\Sigma$ (the LD matrix), where $\Lambda$ is a vector giving the true standardized effect sizes. In a locus with a single causal SNP producing the observed enrichment of signal, and if the causal SNP is in the set modeled by eCAVIAR, $\Lambda$ would consist of a vector with all 0 values except for one SNP with non-zero value. eCAVIAR then models $\Lambda$ with a multivariate normal prior distribution centered on 0 with covariance matrix based on a vector of 0's and 1's giving the true causal status of the SNPs in the locus and a preset scale parameter determined from previous studies.

Here MRLocus diverges from eCAVIAR in two ways. First, we will use a generative model for the elements in the vector $\widehat{\beta}^X_{\cdot,j}$ conditional on the true effect sizes $\beta^X_{\cdot,j}$ within LD-independent signal cluster $j$ (that is, MRLocus models effect sizes rather than $z$-scores). Second, we will use a univariate distribution for each $\widehat{\beta}^X_{i,j}$ instead of a multivariate distribution for the entire vector. This second difference is motivated by practical concerns of model fitting and specification; the univariate modeling provided more efficient model fitting with Stan (higher effective sample sizes and R-hat values near 1), and allowed for more flexible choice of priors, as described below.

Instead of a multivariate normal prior on the estimated coefficients, MRLocus uses a horseshoe prior [Carvalho et al., 2009, 2010], a type of hierarchical shrinkage prior that provides sparsity in the posterior estimates [Piironen and Vehtari, 2017]. In the following, we use the notation of Carvalho et al. [2009], where $\lambda_i$ provides the *local* shrinkage parameters and $\tau$ provides the *global* shrinkage parameter.

The following hierarchical model is fit separately across LD-independent signal clusters $j \in 1, \ldots, J$, and so in the following equations, the subscript for $j$ is omitted for clarity. In all equations below but the last, $i \in 1, \ldots, n_j$. Here, for consistency with Stan code, the normal distribution is written as $N(\mu, \sigma)$ where the second element $\sigma$ provides the standard deviation instead of the variance.

$$\widehat{\beta}^A_i \sim N([\Sigma^A \beta^A]_i, \mathrm{se}(\widehat{\beta}^A_i)) \tag{6}$$
$$\widehat{\beta}^B_i \sim N([\Sigma^B \beta^B]_i, \mathrm{se}(\widehat{\beta}^B_i)) \tag{7}$$
$$\beta^A_i \sim N(0, \lambda_i \tau) \tag{8}$$
$$\beta^B_i \sim N(0, \lambda_i \tau) \tag{9}$$
$$\lambda_i \sim \mathrm{Cauchy}(0, 1) \tag{10}$$
$$\tau \sim \mathrm{Cauchy}(0, 1) \tag{11}$$

Of note, $\beta^A_i$ and $\beta^B_i$ share a prior involving $\lambda_i$, such that evidence from study A and B supporting a SNP $i^\dagger$ that is causal both for eQTL and GWAS signal ($\beta^A_{i^\dagger} \neq 0, \beta^B_{i^\dagger} \neq 0$), will be reflected in larger posterior draws for $\lambda_{i^\dagger}$ compared to other $\lambda_{i'}$ for $i' \neq i^\dagger$.

The posterior mean for $\beta^A_{i,j}$ and $\beta^B_{i,j}$ are the parameters of interest from this step of model fitting, and passed along to the next step after scaling back to the original scale by $1/s^A_j$ and $1/s^B_j$, respectively, as described earlier. We also considered making use of the posterior standard deviation for these two parameters, but found MRLocus gave better performance in terms of accuracy and stability in the Stan fitting procedure if only the posterior mean was kept from the colocalization step. The use of the horseshoe prior in this colocalization step is distinct from other uses of the horseshoe prior in Bayesian Mendelian Randomization methods, Berzuini et al. [2018] and Uche-Ikonne et al. [2019], where it is used as a prior for the pleiotropic effects (effects not mediated by the exposure) or on the mediation slope, respectively.

#### 1.2 Slope fitting step

The second step of MRLocus is to estimate the gene-to-trait effect (the slope in a Mendelian Randomization analysis) using the posterior mean values from the colocalization step. For each LD-independent signal cluster $j$, MRLocus extracts one SNP, based on the largest posterior mean value for $\beta^A_{i,j}$ (prioritizing the putative mediator for SNP selection). MRLocus also has the non-default option to perform model-based clustering using an EM algorithm to determine how many SNPs per LD-independent signal cluster to pass to the slope fitting step [Scrucca et al., 2016], but this option was not extensively evaluated here.

By default, $J$ SNPs are passed from the colocalization step to the slope fitting step, which should represent the candidate causal SNPs from each LD-independent signal cluster. The posterior mean for $\beta^A_{i,j}$ and $\beta^B_{i,j}$ for these SNPs is then provided to the slope fitting step as variables $\widehat{\beta}^A_j$ and $\widehat{\beta}^B_j$. These two variables are modeled as normally distributed variables centered on true values $\beta^A_j$ and $\beta^B_j$ with standard deviation according to the original standard errors $\text{se}(\widehat{\beta}^A_{i,j})$ and $\text{se}(\widehat{\beta}^B_{i,j})$. We found this incorporation of the original standard errors into the slope fitting procedure helped to accurately estimate the uncertainty on the slope (the gene-to-trait effect).

The primary two parameters of interest are the slope $\alpha$, i.e. the predominant gene-to-trait effect demonstrated by the LD-independent signal clusters in the locus, and $\sigma$, the dispersion of gene-to-trait effects from individual signal clusters around the predominant slope. The hierarchical model for the slope fitting step is given by:

$$\widehat{\beta}^A_j \sim N(\beta^A_j, \text{se}^A_j) \tag{12}$$
$$\widehat{\beta}^B_j \sim N(\beta^B_j, \text{se}^B_j) \tag{13}$$
$$\beta^A_j \sim N(0, \text{SD}_\beta) \tag{14}$$
$$\beta^B_j \sim N(\alpha\beta^A_j, \sigma) \tag{15}$$
$$\alpha \sim N(\mu_\alpha, \text{SD}_\alpha) \tag{16}$$
$$\sigma \sim N(0, \text{SD}_\sigma) \tag{17}$$

where the first four equations are defined for $j \in 1, \ldots, J$. In words, $\beta^B_j$ is assumed to follow a normal distribution centered on the predicted value from a line with slope $\alpha$ and no intercept, and with dispersion $\sigma$ around the fitted line. We are both interested in the posterior mean of $\alpha$ and $\sigma$ as well as our uncertainty regarding the point estimates. In particular we focus on a quantile-based credible interval for $\alpha$, which is used for determining our confidence in a gene being a causal mediator for a trait. Large values of $\sigma$ relative to $\alpha$ times typical mediator perturbation sizes ($\beta^A_j$) reflect significant heterogeneity of the gene-to-trait effect.

##### 1.2.1 Choice of hyperparameters

The hyperparameter for the prior for $\beta^A_j$ is $\text{SD}_\beta$, which is set to 2 times the largest absolute value of $\widehat{\beta}^A_j$. The hyperparameters for the prior for $\alpha$ are $\mu_\alpha$ and $\text{SD}_\alpha$, and are set based on a simple un-weighted linear model of $\widehat{\beta}^B_j$ on $\widehat{\beta}^A_j$ without an intercept. $\mu_\alpha$ is set to the estimated slope coefficient, and $\text{SD}_\alpha$ is set to 2 times the absolute value of this estimated slope coefficient.

The hyperparameter for the prior for $\sigma$ is $\text{SD}_\sigma$, which is set by default to 1 (but can be modified by the user). This corresponds to a very wide prior as the GWAS effects sizes are typically quite small in magnitude (as seen in the analyses in the Results).

##### 1.2.2 One signal cluster

In the case that the upstream clumping method only returns a single LD-independent signal cluster, MRLocus relies on a parametric bootstrap to estimate the slope. MRLocus samples the numerator and denominator of the slope from normal distributions $N(\widehat{\beta}_1^B, \text{se}_1^B)$ and $N(\widehat{\beta}_1^A, \text{se}_1^A)$, using the posterior mean estimates from section 1.1. The point estimate is then the median of the bootstrap slope estimates. Intervals are calculated using the MAD of the bootstrap slope estimates, e.g. the median $\pm 1.28$ times the MAD to construct an 80% interval.

### 2 MRLocus Stan code

#### 2.1 Colocalization step

inst/stan/beta_coloc.stan

```
data {
  int n;
  vector[n] beta_hat_a;
  vector[n] beta_hat_b;
  vector[n] se_a;
  vector[n] se_b;
  matrix[n,n] Sigma_a;
  matrix[n,n] Sigma_b;
}
parameters {
  vector[n] beta_a;
  vector[n] beta_b;
  vector<lower=0>[n] lambda;
  real<lower=0> tau;
}
model {
  tau ~ cauchy(0, 1);
  lambda ~ cauchy(0, 1);
  beta_hat_a ~ normal(Sigma_a * beta_a, se_a);
  beta_hat_b ~ normal(Sigma_b * beta_b, se_b);
  for (i in 1:n) {
    beta_a[i] ~ normal(0, lambda[i] * tau);
    beta_b[i] ~ normal(0, lambda[i] * tau);
  }
}
```

#### 2.2 Slope fitting step

inst/stan/slope.stan

```
data {
  int n;
  vector[n] beta_hat_a;
  vector[n] beta_hat_b;
  vector[n] sd_a;
  vector[n] sd_b;
  real sd_beta;
  real mu_alpha;
  real sd_alpha;
  real sd_sigma;
}
parameters {
  real alpha;
  real<lower=0> sigma;
  vector[n] beta_a;
  vector[n] beta_b;
}
model {
  beta_hat_a ~ normal(beta_a, sd_a);
  beta_hat_b ~ normal(beta_b, sd_b);
  beta_a ~ normal(0, sd_beta);
  beta_b ~ normal(alpha * beta_a, sigma);
  alpha ~ normal(mu_alpha, sd_alpha);
  sigma ~ normal(0, sd_sigma);
}
```

#### References


Bob Carpenter, Andrew Gelman, Matthew Hoffman, Daniel Lee, Ben Goodrich, Michael Betancourt, Marcus Brubaker, Jiqiang Guo, Peter Li, and Allen Riddell. Stan: A probabilistic programming language. *Journal of Statistical Software, Articles*, 76(1):1–32, 2017. ISSN 1548-7660. doi: 10.18637/jss.v076.i01.

Stan Development Team. RStan: the R interface to Stan, 2020. URL `http://mc-stan.org/`. R package version 2.21.1.

Pejman Mohammadi, Stephane E. Castel, Andrew A. Brown, and Tuuli Lappalainen. Quantifying the regulatory effect size of cis-acting genetic variation using allelic fold change. *Genome Research*, 27(11):1872–1884, 2017. doi: 10.1101/gr.216747.116.

John Palowitch, Andrey Shabalin, Yi-Hui Zhou, Andrew B. Nobel, and Fred A. Wright. Estimation of cis-eqtl effect sizes using a log of linear model. *Biometrics*, 74(2):616–625, 2018. doi: 10.1111/biom.12810.

Farhad Hormozdiari, Martijn van de Bunt, Ayellet V. Segrè, Xiao Li, Jong Wha J. Joo, Michael Bilow, Jae Hoon Sul, Sriram Sankararaman, Bogdan Pasaniuc, and Eleazar Eskin. Colocalization of gwas and eqtl signals detects target genes. *The American Journal of Human Genetics*, 99(6): 1245–1260, 2020/07/16 2016. doi: 10.1016/j.ajhg.2016.10.003.

Carlos M. Carvalho, Nicholas G. Polson, and James G. Scott. Handling sparsity via the horseshoe. *Proceedings of Machine Learning Research*, 5:73–80, 16–18 Apr 2009. URL `http://proceedings.mlr.press/v5/carvalho09a.html`.

Carlos M. Carvalho, Nicholas G. Polson, and James G. Scott. The horseshoe estimator for sparse signals. *Biometrika*, 97(2):465–480, 04 2010. ISSN 0006-3444. doi: 10.1093/biomet/asq017.

Juho Piironen and Aki Vehtari. On the Hyperprior Choice for the Global Shrinkage Parameter in the Horseshoe Prior. *Proceedings of Machine Learning Research*, 54:905–913, 20–22 Apr 2017. URL `http://proceedings.mlr.press/v54/piironen17a.html`.

Carlo Berzuini, Hui Guo, Stephen Burgess, and Luisa Bernardinelli. A Bayesian approach to Mendelian randomization with multiple pleiotropic variants. *Biostatistics*, 21 (1):86–101, 08 2018. ISSN 1465-4644. doi: 10.1093/biostatistics/kxy027. URL `https://doi.org/10.1093/biostatistics/kxy027`.

Okezie O Uche-Ikonne, Frank Dondelinger, and Tom Palmer. Bayesian estimation of IVW and MR-Egger models for two-sample Mendelian randomization studies. *medRxiv*, 2019. doi: 10.1101/19005868. URL `https://www.medrxiv.org/content/early/2019/09/21/19005868`.

Luca Scrucca, Michael Fop, T. Brendan Murphy, and Adrian E. Raftery. mclust 5: clustering, classification and density estimation using Gaussian finite mixture models. *The R Journal*, 8(1): 289–317, 2016. URL `https://doi.org/10.32614/RJ-2016-021`.
